# Ruxolitinib clears CRYAB p.Arg120Gly aggregates through the ubiquitin-proteasome system

**DOI:** 10.1101/2024.10.11.615348

**Authors:** Erda Alizoti, Leonie Ewald, Simona Parretta, Moritz Meyer-Jens, Ellen Orthey, Christian Conze, Lucie Carrier, Jeffrey Robbins, Sonia R Singh

## Abstract

**Rationale:** Protein accumulation is a hallmark of many neurodegenerative and muscular diseases. Desmin-related (cardio-) myopathy (DRM), a well-studied model for cardiac muscle protein accumulation, is an autosomal dominant-inherited disease presenting with progressive muscle weakness, reduced quality of life, and shortened life span. To date, DRM patients are treated symptomatically and there is no causal treatment available. Independent of the genetic cause, most DRM patients display intracellular accumulation of desmin and its chaperone αB-crystallin (CRYAB). We previously conducted an unbiased high-throughput screen to identify novel effectors that reduce cardiomyocyte aggregate levels and found that downregulation of Janus kinase 1 (JAK1) resulted in lower aggregate load in neonatal mouse cardiomyocytes.

**Objective:** In this study, we tested if the approved JAK inhibitor ruxolitinib ameliorates the disease phenotype in rodent and human CRYAB p.Arg120Gly DRM models.

**Methods and Results:** We found that the mRNA levels of *Jak1* and *Stat3* were higher than any other JAK-signal transducer and activator of transcription (STAT) family members in neonatal rat ventricular myocytes (NRVMs) and human induced pluripotent stem cell-derived cardiomyocytes (hiPSC-CMs). The approved JAK1/2 inhibitor ruxolitinib and the JAK1 inhibitors solcitinib, upadacitinib, and filgotinib prevented accumulation of and cleared pre-existing CRYAB p.Arg120Gly protein aggregates in NRVMs and hiPSC-CMs. Importantly, the knockdown of *Jak1* and *Stat3*, but not *Jak2* resulted in fewer aggregates. Moreover, ruxolitinib, *Jak1* or *Stat3* siRNA treatment enhanced the ubiquitin-proteasome system (UPS)-mediated degradation. Blocking UPS function blunted the effect of ruxolitinib or *Jak1* siRNA on CRYAB p.Arg120Gly accumulation. RNAseq of NRVMs treated with *Jak1* siRNA extracts revealed higher gene expression of important muscle E3 ubiquitinating enzymes. Knockdown of the E3 ligase *Asb2* (Ankyrin Repeat And SOCS Box Protein 2) abolished the effect of ruxolitinib on CRYAB p.Arg120Gly aggregates. In DRM mice, phospho-STAT3 levels were markedly higher than in non-transgenic (NTG) mice with age. Ruxolitinib treatment or *Jak1* knockout prevented cardiac dysfunction and reduced CRYAB p.Arg120Gly aggregate load in DRM mice.

**Conclusion:** In this study, we uncovered the previously unknown effect of the approved drug ruxolitinib to enhance UPS-mediated degradation and prevent protein aggregates in cardiomyocytes.

## Introduction

Intracellular protein accumulations are particularly toxic to non-dividing cells with high energy demand such as (cardio-) myocytes or neurons and result in severe myopathies or neurodegenerative diseases.^1^ Genetic variants in desmin (*DES*), αB-crystallin (*CRYAB*), myotilin (*MYOT*), Z-band alternatively spliced PDZ-motif (*ZASP*), filamin C (*FLNC*), BCL2- associated athanogene 3 (*BAG3*), and four and a half LIM domain protein (*FHL1*) cause desmin-related (cardio-) myopathy (DRM) with intracellular protein accumulations. To date, DRM patients are treated symptomatically, and no causal treatment is available. Different types of cardiomyopathy have been associated with DRM, such as dilated, hypertrophic, restrictive, and arrhythmogenic cardiomyopathies.^2^

Independent of the genetic cause and type of cardiomyopathy, most DRM patients display intracellular accumulation of desmin and its chaperone αB-crystallin (CRYAB). Desmin is a part of the type III intermediate filament protein group, interacting with myofibrils at the Z-disc, therefore building a cellular scaffold that maintains a spatial relationship between the contractile apparatus and other components of the cardiomyocyte. Thus, its main functions include preserving cellular integrity, force transmission, and mechanochemical signaling.^3, 4^ CRYAB is a small heat shock protein and chaperone that builds oligomeric complexes, interacts with desmin to support its proper folding, and prevents its aggregation.^5^

The *CRYAB* c.358A>C (p.Arg120Gly; R120G) variant was first identified in a French family^6^ and has been intensively studied in rodent models.^7–10^ Mouse or rat cardiomyocytes overexpressing CRYAB^R120G^ present large intracellular aggregates, disrupted cellular organization, and mitochondrial and proteolytic system impairment.^11^ CRYAB^R120G^ mice develop cardiac hypertrophy and dilation, reduced ejection fraction with age, and die around 7-8 months.^7, 9^ We previously conducted a high throughput screen to identify novel effectors that prevent CRYAB p.Arg120Gly protein aggregation in neonatal mouse cardiomyocytes,^12^ and we identified Janus kinase 1 (JAK1) as a novel target in protein aggregation. JAK1 belongs with JAK2, JAK3, and TYK2 to the family of non-receptor tyrosine kinases. The canonical function of JAKs is to transmit extracellular signals such as cytokines binding to a receptor tyrosine kinase, via phosphorylation of Signal Transducers and Activators of Transcription (STATs), which then translocate into the nucleus to regulate gene expression.^13^ The JAK-STAT pathway regulates cell proliferation, differentiation, and survival and is especially important for immune function and hematopoiesis.^13^ Thus, JAK variants can cause a wide range of diseases from malignancies to autoimmune diseases.^14^ Therefore, several JAK inhibitors (Jakinibs) have been identified and tested in clinical trials recently with ruxolitinib (Jakafi^TM^), being the first FDA-approved Jakinib drug.

In this study, we investigated the effect of ruxolitinib and other Jakinibs on CRYAB p.Arg120Gly aggregation in neonatal rat ventricular myocytes (NRVMs) and human induced pluripotent stem cell-derived cardiomyocytes (hiPSC-CMs), the effect of ruxolitinib on the function of proteolytic systems, and alterations in gene expression by JAK1 loss-of-function, as well as STAT3 phosphorylation levels, short-term ruxolitinib treatment, and *Jak1* knockout in CRYAB p.Arg120Gly mice with DRM.

## Materials and Methods

### Isolation of neonatal rat ventricular myocytes (NRVMs)

Isolation of hearts from neonatal rats was performed by decapitation without anesthetic agents and conformed to the guidelines from directive 2010/63/EU of the European parliament on the protection of animals used for scientific purposes and the corresponding German authorities (Hamburg, ORG949; for isolations done in Germany) and the NIH Guide for Care and Use of Laboratory Animals (for isolations done in the USA). Hearts were harvested from 1-3-day-old pups (Sprague Dawley or Wistar), washed in Calcium and Bicarbonate-free Hanks with HEPES (CBFHH) buffer, cut into small pieces and dissociated 3-5 min step-wise with 0.25% trypsin (Difco, 0152-15-9) at RT. Dissociated cells were collected in DNase (Sigma, D8764)/FCS (active) solution, spun down at 60 g and 4°C for 15 min, resuspended in medium (DMEM, 10% FCS, 1% P/S) and dropped through a cell strain. Cells were then pre-plated for 60-90 min to reduce the number of non-cardiomyocytes in the supernatant.

### Human induced pluripotent stem cell (hiPSC) culture and cardiomyocyte (hiPSC-CM) differentiation

UKEi001-A^15^ (ERC001) or mTagRFPT-TUBA1B (AICS-0031-035, Coriell Institute; for JAK-STAT pathway expression values) hiPSCs were expanded and differentiated using either embryoid body and growth factor-based three stage protocol^16^ or a small-molecule based protocol according to Pietsch et al.^17^ In brief, the expansion of the hiPSCs was performed in FTDA medium (house-made, DMEM F-12, L-glutamine 2 mM, transferrin 5 μg/ml, selenium 5 ng/ml, human serum albumin 0.1 %, lipid mixture 1x, insulin 5 μg/ml, dorsomorphin 50 nM, activin A 2.5 ng/ml, TGFβ1 0.5 ng/ml, bFGF 30 ng/ml) on Geltrex^TM^-coated (Gibco A14133– 02) cell culture vessels. Cardiomyocyte differentiation was performed by treating high-density undifferentiated monolayer iPSC cultures with an activin A/bone morphogenic factor (BMP4) based protocol. After dissociation, hiPSC-CMs were seeded at a density of 5 x 10^6^ cells/well in complete medium (house-made, DMEM, penicillin-streptomycin 1%, horse serum (inactive) 10%, insulin 10 μg/ml, aprotinin 33 μg/ml) on Geltrex^TM^ (Gibco A14133–02)-coated 6-well culture plates (Nalge Nunc International, 154461) and kept at 37°C and 7% CO_2_ with medium change on Mondays, Wednesdays and Fridays.

### siRNA transfection, compound treatment and virus transduction of cardiomyocytes

NRVMs were seeded into 0.1% gelatine-coated (Sigma-Aldrich, G1393) 12- or 6-well plates, Lab-Teks (Nalge Nunc International, 154,461) or 6-well plates with nanopattern coverslips (Tebu-Bio ANFS-CS25 904). Per 12-, 6-well plate, or LabTek well, 1.5 x 10^5^, 3 x 10^5^, or 2 x 10^5^ cells were seeded, respectively. HiPSC-CMs were seeded on Geltrex^TM^-coated 6-well plates with nanopattern coverslips or 96-well plates (ThermoFisher Scientific, 165305). Per 6-well plate or 96-well plate well, 3 x 10^5^, or 1 x 10^4^ cells were seeded, respectively. For siRNA transfection of NRVMs, the medium was changed to 500 µL (12-well and LabTeks) or 1 mL (6-well) OptiMEM (Thermo Fisher Scientific, 51985–026). SiRNAs and Lipofectamine3000 (Thermo Fisher Scientific, L3000008) transfection reagent (final amount of 4 µL/mL) were diluted in 50 µL (12-well and LabTeks) or 100 µL (6-well) OptiMEM each. After 5 minutes of incubation, the solutions were mixed and incubated for 20 minutes before adding them dropwise to the wells. Four hours later, DMEM with 20% FBS was added vol/vol, or cells were transduced with adenovirus. Two days later, the medium was changed to culture medium with 10 µM cytosine β-D-arabinofuranoside (Sigma-Aldrich, C1768).

Following siRNAs were used as indicated: scramble siRNA (corresponding concentration, Thermo Fisher Scientific, 4390846), siJak1 (Thermo Fisher Scientific, s7646-48), siJak2 (Thermo Fisher Scientific, s7650), siStat3 (Thermo Fisher Scientific, s129048), siPsmd1 (20 nM; Thermo Fisher Scientific, s136286), siAsb2 (Thermo Fisher Scientific, s28515), siRapsn (Thermo Fisher Scientific, s168618), siRnf207 (Thermo Fisher Scientific, s193639), or siTrim50 (Thermo Fisher Scientific, s143615).

For overexpression of CRYAB^R120G^-GFP, GFP, and GFPu, NRVMs or hiPSC-CMs were transduced with an adenovirus serotype 5 (AdV5) with CMV promoter. The appropriate virus amount (MOI 0.5-10) was determined in preliminary experiments and varied between virus productions. Of note, hiPSC-CMs were transduced with about double the amount of virus than NRVMs. For transduction, the virus was diluted in 500 µL (12-well and LabTeks) or 1 mL (6-well) OptiMEM, and the medium was replaced by the solution for 2 hours. If cells were transfected before transduction, the virus was diluted in 250 µL (12-well and LabTeks) or 500 µL (6-well) OptiMEM and added to the transfection solution after 4 hours. After 2 hours of virus transduction, DMEM with 20% FBS (NRVMs) or double concentrated hiPSC-CM culture medium was added vol/vol.

All compounds were dissolved in dimethylsulfoxide (DMSO). The final concentration of DMSO was 0.1%. The JAK inhibitors ruxolitinib (3 µM), filgotinib (1 µM), solcitinib (1 µM), and upadacitinib (1 µM) were added either from day 1 with the double-concentrated medium) or as indicated. For live cell imaging, 10 µM ruxolitinib or 0.1% DMSO was added once right before the experiment. The culture medium with inhibitors was refreshed every other day.

For autophagic flux measurement, cells were treated with 50 nM bafilomycin A_1_ (Sigma-Aldrich, B1793) and/or 50 nM torin 2 (LC laboratories, T8448) for 3 hours.

### Mouse experiments

The investigation conformed to the guide for the care and use of laboratory animals published by the National Institutes of Health (Publication No. 85-23, revised 2011, published by the National Research Council). Cardiomyocyte-specific (*Myh6* promoter) transgenic CRYAB p.Arg120Gly^7^ were maintained on FVBN background. Conditional *Jak1* knockout (KO) C57BL/6 mice were purchased from Taconic Biosciences (C57BL/6-Jak1^tm1.1^ ^mrl^) and crossed with CRYAB p.Arg120Gly and cardiomyocyte-specific (*Myh6* promoter) MerCreMer (MCM) mice^18^ on FVBN background for at least five generations. Mice were sacrificed by cervical dislocation under isoflurane or CO_2_-anesthesia. After thoracotomy, hearts were extracted, rinsed in sodium chloride, and shortly dried on a paper tissue. Organs were fixed in formalin or ROTI® Histofix (Carl Roth, P087.1), or shock frozen in liquid nitrogen.

A Visual Sonics Vevo 2100 Imaging System with a 40 MHz transducer was used for two-dimensional guided M-mode echocardiography following isoflurane anesthesia (performed blinded by Cincinnati Children’s Hospital Heart Institute Research Core).

Ruxolitinib was dissolved in DMSO (100 µg/µL) and diluted in water containing 0.5% methylcellulose (w/v) and 0.1% Tween 80 for administration of 75 mg/kg ruxolitinib twice daily by oral gavage (according to Heine et al.^19^). The vehicle group was treated with corresponding amount of DMSO in water containing 0.5% methylcellulose (w/v) and 0.1% Tween 80. Conditional *Jak1* KO was induced in 8-week-old mice with tamoxifen-supplemented chow according to Meng et al.^20^.

### RNA extraction, RT-qPCR, RNA sequencing, and NanoString RNA analysis

For the measurement of gene expression levels, NRVMs were lysed in 500 µL (12-well) RNAzol RT Reagent (Molecular Research Center Inc, RN 190). For the measurement of gene expression levels, NRVMs or powdered mouse hearts were lysed in 500 µL (12-well) or 1 mL (30 mg heart powder) RNAzol RT Reagent (Molecular Research Center Inc, RN 190). CDNA synthesis was performed with iScript cDNA synthesis kit (Bio-Rad, 1,708,840) with 100-200 ng RNA. RT-qPCR was then performed with TaqMan gene expression assays (Thermo Fisher Scientific; Psmd1 - Rn01400483_m1, Jak1 - Mm00600614_m1, Jak2 -Mm01208489_m1, Stat1 - Mm01257286_m1, Stat3 - Mm01219775_m1, Tyk2 -Mm00444469_m1, 18s - Hs03003631_g1) and SsoAdvancedTM Universal Probes Supermix (BioRad #1725281). RNA sequencing was performed as described before^21^ by the Cincinnati Children’s Hospital and Medical Center DNA core with RNA polyA-stranded library preparation, paired-end 75 bp sequencing conditions and 20 M reads per sample. Three samples (technical replicates) were used from 3 preparations (biological replicates).

NanoString RNA analysis was performed and analyzed by the UKE NanoString Facility using nCounter Gene Expression Assay.

### Protein extraction and Western blot

For Western blot analysis, cells were harvested in CelLytic^TM^ M (Sigma-Aldrich, C2978, soluble fraction) or a SDS-based lysis buffer (3% SDS, 30 mM Tris base, pH 8.8, 5 mM EDTA, 30 mM NaF, 10% glycerol, 1 mM DTT, crude or sarcomere- and membrane-enriched fraction) with protease inhibitor cocktail (cOmplete^TM^ mini, EDTA-free, Roche, 11836170001 and PhosSTOP^TM^, Roche, PHOSS-RO). Mouse tissue samples were powdered and about 30 mg of powder were dissolved in 180 μL CelLytic M with protease inhibitor cocktail and homogenized by using a tissue lyzer (2 x 30 s at 20 Hz) and centrifuged at 4 °C, full speed for 30 min in a table-top centrifuge. The pellet of the first step was homogenized in 180 μL SDS buffer and centrifuged at room temperature, full speed for 15 min in a table-top centrifuge. Afterward, 8–25 μg protein with 6x Laemmli buffer was loaded onto 4-15% or 4– 20% Mini-PROTEAN TGX^TM^ Precast Protein Gels (Bio-Rad, 4561084) or 8 or 12% self-casted acrylamide/bisacrylamide (29:1) gels. After electrophoresis, the proteins were transferred to nitrocellulose or PVDF membranes with Bio-Rad Tris/Glycine buffer or self-made transfer buffer (25 mM Tris base, 190 mM glycine, 20% methanol, pH 8.3) and stained with the following antibodies: JAK1 – R&D systems, MAB4260; JAK2 – Cell Signaling Technology, 3230; STAT3 - Cell Signaling Technology, 12640; P-STAT3 – Cell Signaling Technology, 9145S; GAPDH – HyTest, 5G4 or Sigma-Aldrich, P0067; ACTN2 - Sigma-Aldrich, A7811; GFP – Santa Cruz Biotechnology, sc-9996; LC3B – Cell Signaling Technology, 2775; p70 S6K – Cell Signaling, 34475. Western blots were either detected with LI-COR Biosciences Odyssey, Biorad ChemiDoc, or Vilber Fusion FX7. Quantification was done with Image Studio (LICOR), Image Lab (Bio-Rad), or Bio-1D (Vilber) software.

### Fluorescence imaging

NRVMs or hiPSC-CMs were fixed in 4% PFA for 15-20 mins at 4°C, permeabilized for 15 mins in PBS/0.5% Triton X100, and stained for the cardiomyocyte marker TNNI3 (troponin I, cardiac 3; Millipore, MAB1691) or ACTN2 (Sigma-Aldrich, A7811), nuclei (DAPI, Thermo Fisher Scientific, D1306) or phalloidin (phalloidin-Alexa Fluor 633 Thermo Fisher Scientific A22284) in blocking solution (PBS, 1% BSA, 0.1% Tween 20, 0.05% sodium azide) and mounted with ProLongTM Gold antifade (Thermo Fisher Scientific #P36930) for LabTeks or nanopatterned cover slips or kept (well plates) in PBS for imaging. Mouse hearts were fixed in formalin or ROTI® Histofix (Carl Roth, P087.1), embedded in paraffin, sliced and stained with for the cardiomyocyte marker TNNI3 (Millipore, MAB1691), nuclei (DAPI) and CRYAB (Stressgen, SPA-222). Slides or plates were imaged using a Nikon Eclipse Ti inverted microscope equipped with an Andor Zyla-4.2 sCMOS camera and using a Plan Apo λ 20x/0.75 objective lens or a Zeiss Axio Observer.Z1/7 microscope LSM800 with a Plan-Apochromat 20x/0.80 objective lens. For live cell imaging, cells were stained with SiR-actin (Spirochrome, SC001) or SPY-tubulin (Spirochrome, SC503) and imaged with a Nikon Eclipse Ti2-E inverted microscope equipped with an Okolab incubation chamber or Nikon BioStation IM-Q time-lapse imaging system with a 40x/0.80 objective lens. Whole images were quantified for aggregates within cardiomyocytes or F-actin-positive cells with NIS Elements software or ImageJ by setting a threshold for each channel, using the default method, and defining the cardiomyocyte area as a region of interest (ROI).

### Statistical analysis

Statistical tests were performed with GraphPad Prism 8. The applied test and p-value are indicated in the figure or figure legend. A p-value > 0.05 was considered significant.

## Results

*Ruxolitinib treatment prevents CRYAB p.Arg120Gly aggregate formation and partially clears pre-existing aggregates in NRVMs*

RNA-seq of NRVMs and hiPSC-CMs revealed that *Jak1*/*JAK1* and *Stat3*/*STAT3* are the most abundant of JAK-STAT family members, suggesting their important role in cardiomyocytes (Table S1). Using different concentrations of the JAK inhibitor ruxolitinib in NRVMs, we showed that 3 µM ruxolitinib was sufficient to abolish phosphorylated STAT3 protein levels (Figure S1). To test if ruxolitinib treatment affected CRYAB p.Arg120Gly aggregate load, NRVMs were transduced with AdV5-CRYAB^R120G^-GFP and kept in culture for 10 days (Figure 1). Since DRM patients already show aggregate accumulation in muscle biopsies at the time they seek health care, it is of great interest that a potential drug can prevent further protein aggregation at a late stage to blunt disease progression and ideally clear pre-existing aggregates to potentially heal from the disease. To determine whether ruxolitnib could i) prevent aggregate formation at an early stage, ii) prevent aggregate formation at a late stage and iii) clear pre-existing aggregates, ruxolitnib was added either from day 1, 3, 5, or 7. The aggregates of CRYAB p.Arg120Gly grew over time (Figure 1A images left panel; Figure 1B quantification purple line). Ruxolitinib treatment from day 1 very efficiently prevented CRYAB p.Arg120Gly aggregate formation (Figure 1A images second panel; Figure 1B quantification black line). Adding ruxolitinib at later time points (day 3, 5, or 7) prevented further formation of aggregates and ruxolitinib treatment from day 3 or 5, indicating that pre-existing aggregates could be cleared (Figure 1A images 3^rd^-5^th^ panel; Figure 1B quantification blue, orange and yellow lines). Ruxolitinib did not result in lower GFP levels in a control experiment (Figure S2A and B).

**Figure 1.**
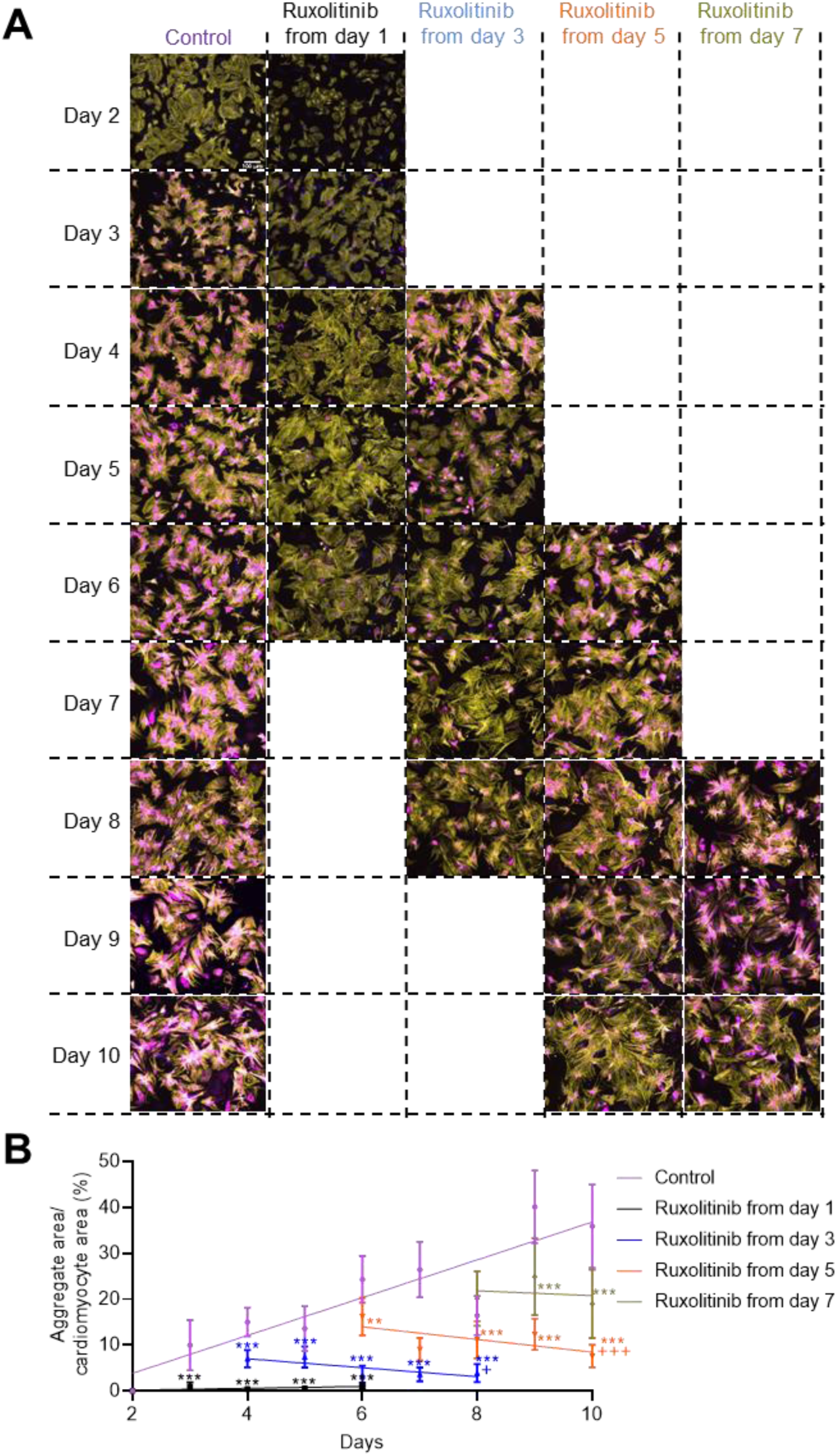
Ruxolitinib treatment prevents CRYAB p.Arg120Gly aggregate formation at early and late stages and partially clears pre-existing aggregates. NRVMs were transduced with AdV5-CMV- CRYAB^R120G^-GFP and treated with either 3 µM ruxolitinib or DMSO on the indicated day. Medium change with ruxolitinib or DMSO was performed every other day. **A,** Representative IF images. Aggregates are depicted in magenta (CRYAB^R120G^-GFP), cardiomyocytes in yellow (anti-cardiac troponin I), and nuclei in blue (DAPI). Scale bar = 100 µm. **B,** Quantification of aggregates in cardiomyocytes with NIS Elements software. Data were obtained from 2 independent NRVM preparations (prep) with 3 wells per prep, 7-8 images per well and are depicted as mean ± SEM, one-way ANOVA, Dunnett’s post-hoc analysis for same time points with **p<0.01 and ***p<0.001 or unpaired Student’s t-test for comparison of start and end of the experiment within one group with ^+^p<0.05 and ^+++^p<0.001.

### Ruxolitinib treatment dissolves CRYAB p.Arg120Gly aggregates in NRVMs

To visualize and confirm that ruxolitinib cleared pre-existing CRYAB p.Arg120Gly aggregates, we performed live cell imaging in NRVMs transduced with CRYAB^R120G^-GFP. Ruxolitinib (10 µM) was added at the beginning of the experiment and the compound was not renewed due to technical limitations. As expected, ruxolitinib treatment led to fewer CRYAB p.Arg120Gly aggregates after 2.5 days (endpoint; Figure 2A-C). Strikingly, some of the ruxolitinib-treated cells exhibited aggregate dissolutions (Figure 2A and B; marked with arrow). However, not all cells in the ruxolitinib-treated samples showed that dissolution.

**Figure 2.**
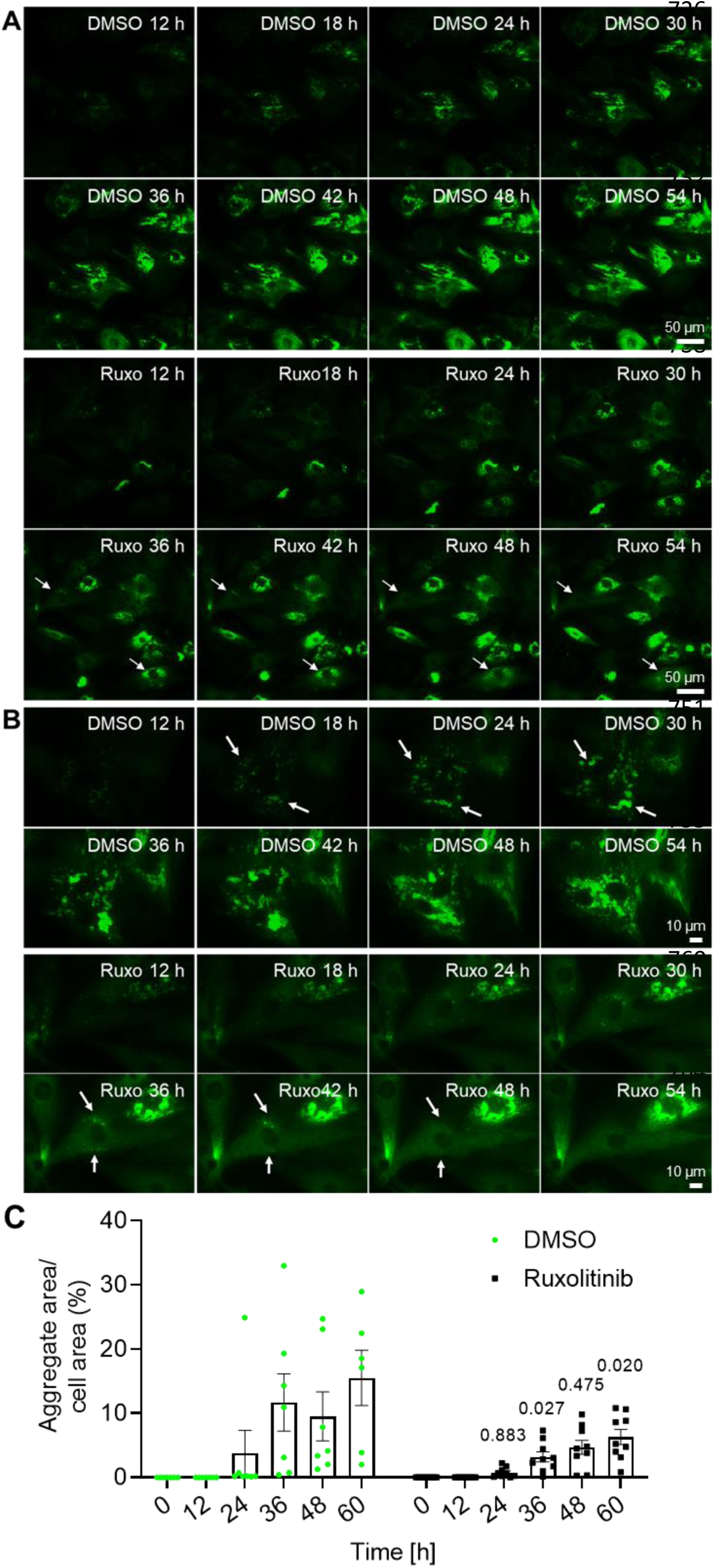
Ruxolitinib dissolves CRYAB p.Arg120Gly aggregates in NRVMs. Live cell imaging of NRVMs transduced with Ad5-CMV-CRYAB^R120G^-GFP and treated with 10 µM ruxolitinib (ruxo) or DMSO. **A**, Representative fluorescence images. Scale bar = 50 µm. **B**, Zoom of representative cardiomyocytes. Aggregates are depicted in green (CRYAB^R120G^-GFP). Arrows mark growing or dissolving aggregates. Scale bar = 10 µm. **C**, Quantification of aggregates in actin area (not shown; SiR-actin staining) with NIS Elements software. Each data point represents a whole image with a number of cells. Data were obtained from 8-10 points from 2 wells per condition and 1 NRVM preparation and are depicted as mean ± SEM, and p-values were obtained with two-way ANOVA and Sidak’s multiple comparisons post-hoc analysis. Corresponding videos can be found in the online supplements.

### JAK1 inhibitors result in fewer CRYAB p.Arg120Gly aggregates in human cardiomyocytes

To test if ruxolitinib is as effective on CRYAB p.Arg120Gly aggregates in human cardiomyocytes, we performed a live cell imaging experiment in hiPSC-CMs transduced with CRYAB^R120G^-GFP and found that ruxolitinib there also cleared pre-existing aggregates in a portion of cells (Figure 3A; indicated by arrows). The video quantification confirmed fewer CRYAB p.Arg120Gly aggregates in ruxolitinib-treated hiPSC-CMs at 36, 48, and 60 h (Figure 3B). We then tested the more specific JAK1 inhibitors solcitinib, filgotinib (Jyseleca^TM^), and upadacitinib (Rinvoq^TM^) in hiPSC-CMs. Filgotinib and upadacitinib are approved for the treatment of rheumatoid arthritis and ulcerative colitis. All three inhibitors are reversible ATP-binding site inhibitors with at least 5-fold higher selectivity for JAK1 than for JAK2, JAK2, or TYK2. HiPSC-CMs were transduced with AdV5-CRYAB^R120G^-GFP and treated with ruxolitinib, solcitinib, upadacitinib, filgotinib, or DMSO (vehicle control; Figure 3C). All tested compounds resulted in lower CRYAB p.Arg120Gly aggregate load in hiPSC-CMs (Figure 3C and D). A similar result was obtained with solcitinib, upadacitinib, and filgotinib in NRVMs (Figure S4). All inhibitors reduced CRYAB p.Arg120Gly aggregate load by at least 50%. Thus, JAK inhibition reduces CRYAB p.Arg120Gly aggregate load also in human cardiomyocytes.

**Figure 3.**
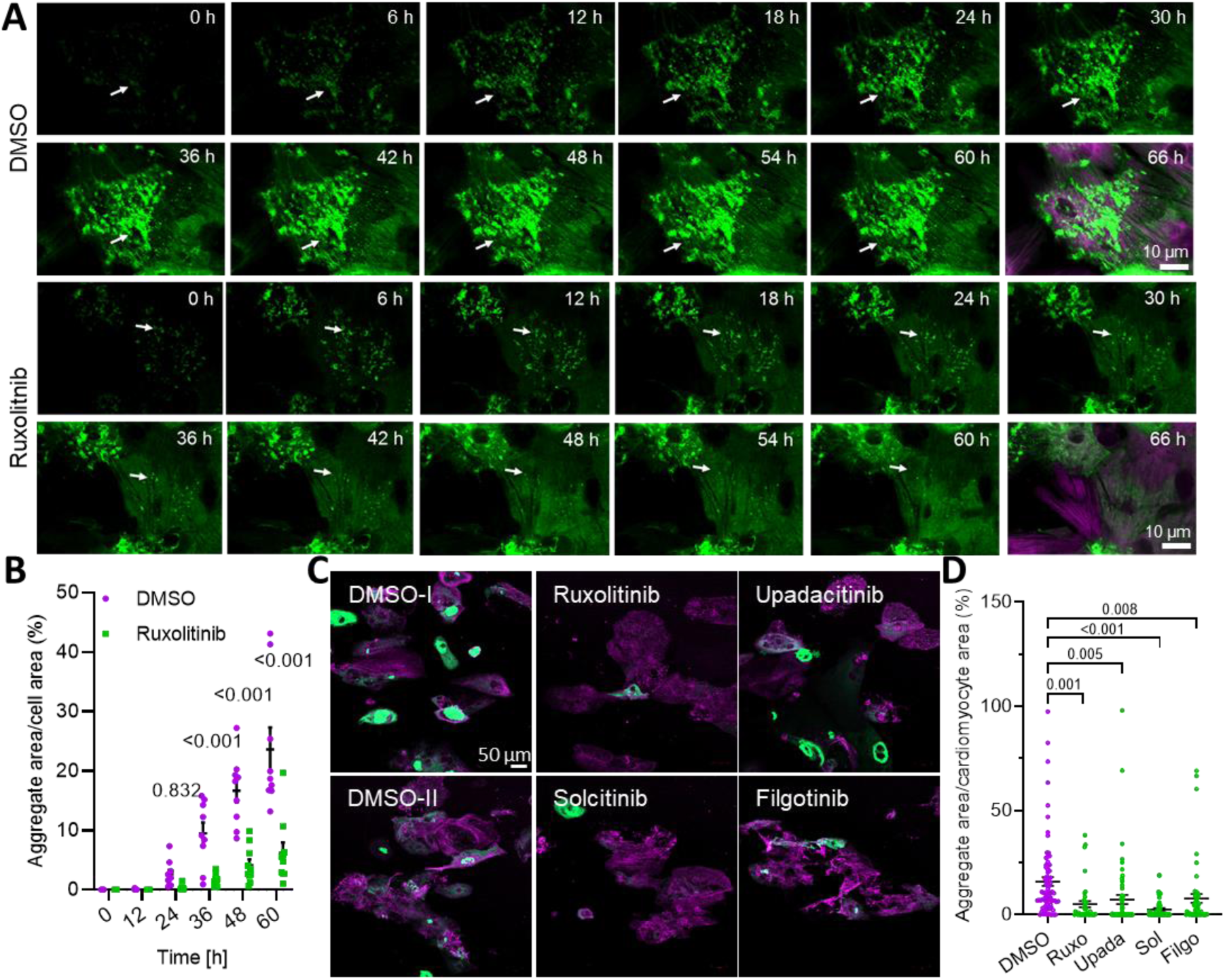
JAK1 inhibitors result in fewer CRYAB p.Arg120Gly aggregates in human cardiomyocytes. **A**, HiPSC-CMs transduced with AdV5-CMV-CRYAB^R120G^-GFP were treated with 10 µM ruxolitinib or 0.1% DMSO at the start of the long-term time-lapse live cell imaging. Representative images. Scale bar = 10 µm. Aggregates are depicted in green (CRYAB^R120G^-GFP) and cells in magenta (SiR-actin). Arrows indicate moving and forming, or dissolving aggregates. The corresponding videos can be found in the supplements. **B**, Quantification of aggregates in F-actin-positive cells of videos from A with NIS Elements software. Data were obtained from one hiPSC-CM differentiation with two wells per condition and 5-8 selected points per well and are depicted as mean ± SEM, with p-values from two-way ANOVA and Sidak’s post-hoc analysis. **C**, Representative IF images of fixed hiPSC-CMs transduced with AdV5-CMV-CRYAB^R120G^-GFP and treated with 3 µM ruxolitinib (Ruxo), 1 µM upadacitinib (Upada), 1 µM solcitinib (Sol), 1 µM filgotinib (Filgo) or 0.1% DMSO every other day for 1 week. Aggregates are depicted in green (CRYAB^R120G^-GFP and cardiomyocytes in magenta (anti-ACTN2). Scale bar = 50 µm. **D**, Quantification of aggregates in cardiomyocytes (hiPSC-CMs) from B with ImageJ software. Data were obtained from 2 independent hiPSC-CM differentiations with at least 4 wells per condition and 5-8 images per well, and are depicted as mean ± SEM, with p-values obtained by the one-way ANOVA and Dunnett’s post-hoc analysis.

### Knockdown of Jak1 and Stat3 but not Jak2 prevents CRYAB p.Arg120Gly aggregate formation

We further wanted to test whether the prevention of aggregate formation is caused by the inhibition of JAK1 and if this effect is mediated through a STAT or potentially a non-canonical JAK function. Thus, we tested siRNAs targeting *Jak1*, *Jak2,* or *Stat3* in NRVMs. Ruxolitinib treatment itself did not affect protein levels of JAK1 (Figure 4A), JAK2, or STAT3, but abolished STAT3 phosphorylation (Figure 4A; Figure S2B). As expected, NRVMs transduced with AdV5-CRYAB^R120G^-GFP and treated with ruxolitinib developed very few CRYAB p.Arg120Gly aggregates (Figure 4A). JAK1 protein level was 60% lower in NRVMs treated with siJak1 than with scramble siRNA (scr; Figure 4B). This was associated with markedly lower CRYAB p.Arg120Gly aggregate load, suggesting that a partial reduction of JAK1 function is sufficient to prevent aggregate formation. *Jak1* knockdown did not lead to lower GFP levels in a control experiment (Figure S2C). The level of JAK2 protein was markedly lower in cells treated with siJak2 than with scr, but did not affect the abundance of CRYAB p.Arg120Gly aggregates (Figure 4C), suggesting that protein accumulation is not regulated by JAK2. In addition, we tested if canonical JAK1-STAT3 signaling is responsible for the observed effects on the aggregates. STAT3 protein level and CRYAB p.Arg120Gly aggregate load were both markedly lower in NRVMs treated with siStat3 than with scr (Figure 4D), suggesting that reduction of CRYAB p.Arg120Gly aggregates is mediated through JAK1-STAT3.

**Figure 4.**
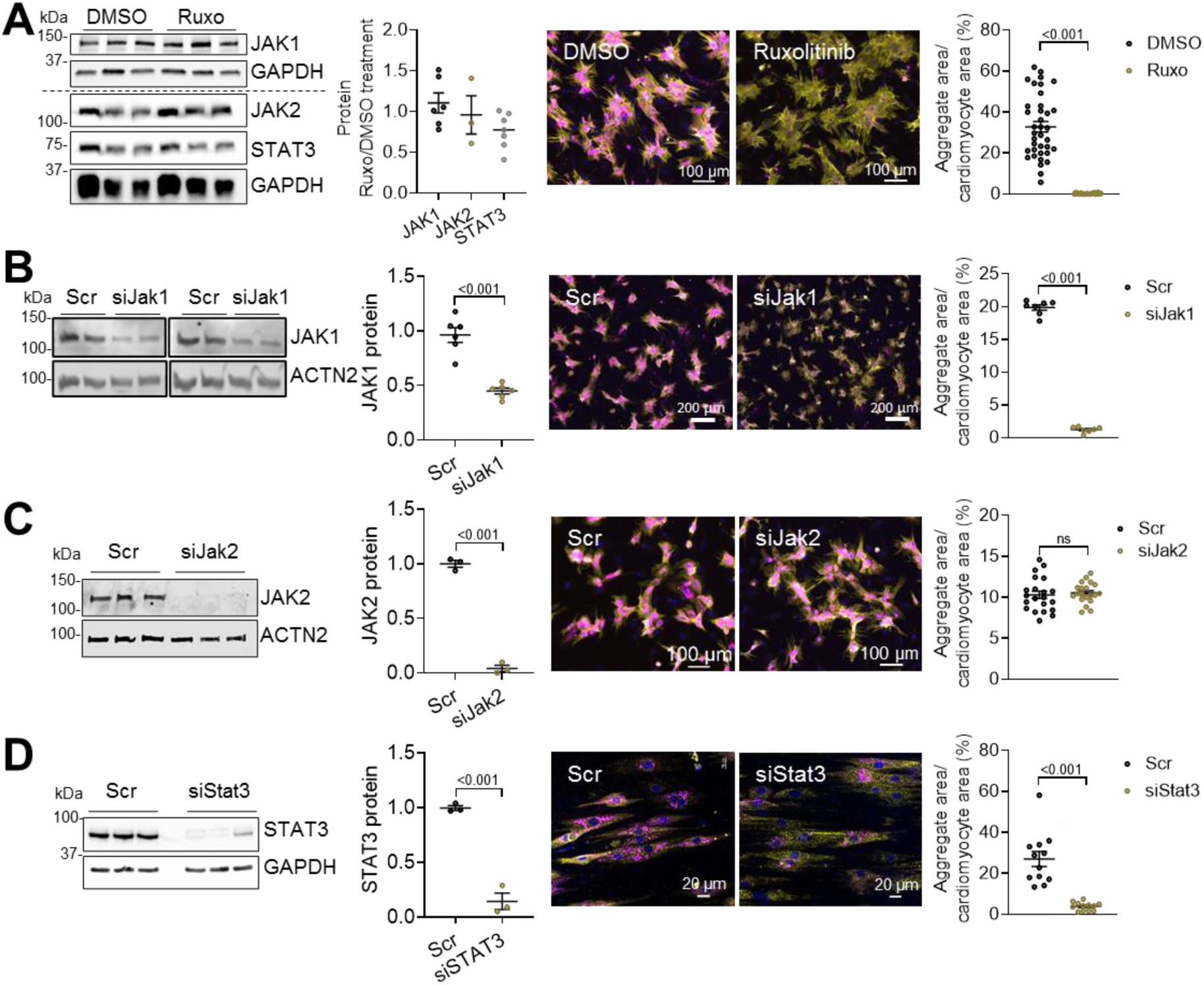
Knockdown of *Jak1* or *Stat3* but not *Jak2* results in markedly lower CRYAB p.Arg120Gly aggregate load. NRVMs were transduced with AdV5-CMV-CRYAB^R120G^-GFP and treated with 3 µM ruxolitinib or DMSO or 100 nM siRNAs targeting either JAK1 (siJak1), JAK2 (siJak2), STAT3 (siStat3) or scramble siRNA (scr). Thereafter, NRVMs were harvested or fixed after 4-6 days. **A**, Treatment with ruxolitinib or DMSO; Scale bar = 100 µm. **B**, Transfection with siJak1; Scale bar = 200 µm. **C**, Transfection with siJak2; Scale bar = 100 µm **D**, Transfection with siStat3, Scale bar = 20 µm. **A**, **B**, **C**, **D**, Western blots of protein extracts from treated NRVMs were stained with antibodies directed against indicated proteins. In the representative immunofluorescence images, aggregates are depicted in magenta (CRYAB^R120G^-GFP), cardiomyocytes in yellow (anti-cardiac troponin I), and nuclei in blue (DAPI). Quantification of aggregates in cardiomyocytes with NIS Elements or ImageJ software. Data were obtained from 1 (Western blot) or 2 (immunofluorescence) NRVM preparations with at least 3 wells per condition and at least 7 images per well for immunofluorescence. Data are depicted as mean ± SEM, and p-values were obtained with the unpaired Student’s t-test. Abbreviation: ns, non-significant.

### UPS- but not ALP-mediated degradation is enhanced by ruxolitinib treatment, Jak1 or Stat3 knockdown

Since proteolytic pathways such as the UPS or autophagy-lysosomal pathway (ALP) play a major role in the removal of protein accumulations and aggregates, we tested the effect of ruxolitinib, siJak1 or siStat3 treatment on UPS- or ALP-mediated degradation. To our knowledge, the UPS- or ALP-mediated degradation regulation of ruxolitinib has not yet been reported. To assess the UPS-mediated degradation, NRVMs were transduced with an AdV5- GFPu (inverse UPS reporter), transfected with siPsmd1 (knockdown of UPS function) or scr, and treated with ruxolitinib or DMSO (Figure 5A). As expected,^21^ efficient knockdown of *Psmd1*, encoding the essential 26S proteasome non-ATPase regulatory subunit 1 (PSMD1) led to higher GFPu protein levels (Figure 5A), suggesting an impaired UPS-mediated degradation. In line with previous findings,^22^ transduction of NRVMs with AdV5-CRYAB^R120G^- GFP also impaired UPS-mediated degradation (Figure 5A). Ruxolitinib treatment resulted in markedly lower GFPu protein levels in CRYAB^R120G^-GFP-transduced cells, suggesting that it positively regulates UPS-mediated degradation (Figure 5A). This effect of ruxolitinib was abolished when NRVMs were treated with siPsmd1 (Figure 5A). In line with these results, siJak1 treatment resulted in a markedly lower GFPu protein level, which was blunted with *Psmd1* knockdown (Figure 5B). Similarly, *Stat3* knockdown led to lower GFPu protein levels (Figure 5C). In summary, these experiments suggest a positive regulation of UPS-mediated degradation by ruxolitinib, mediated through a loss of JAK1 and STAT3 function.

**Figure 5.**
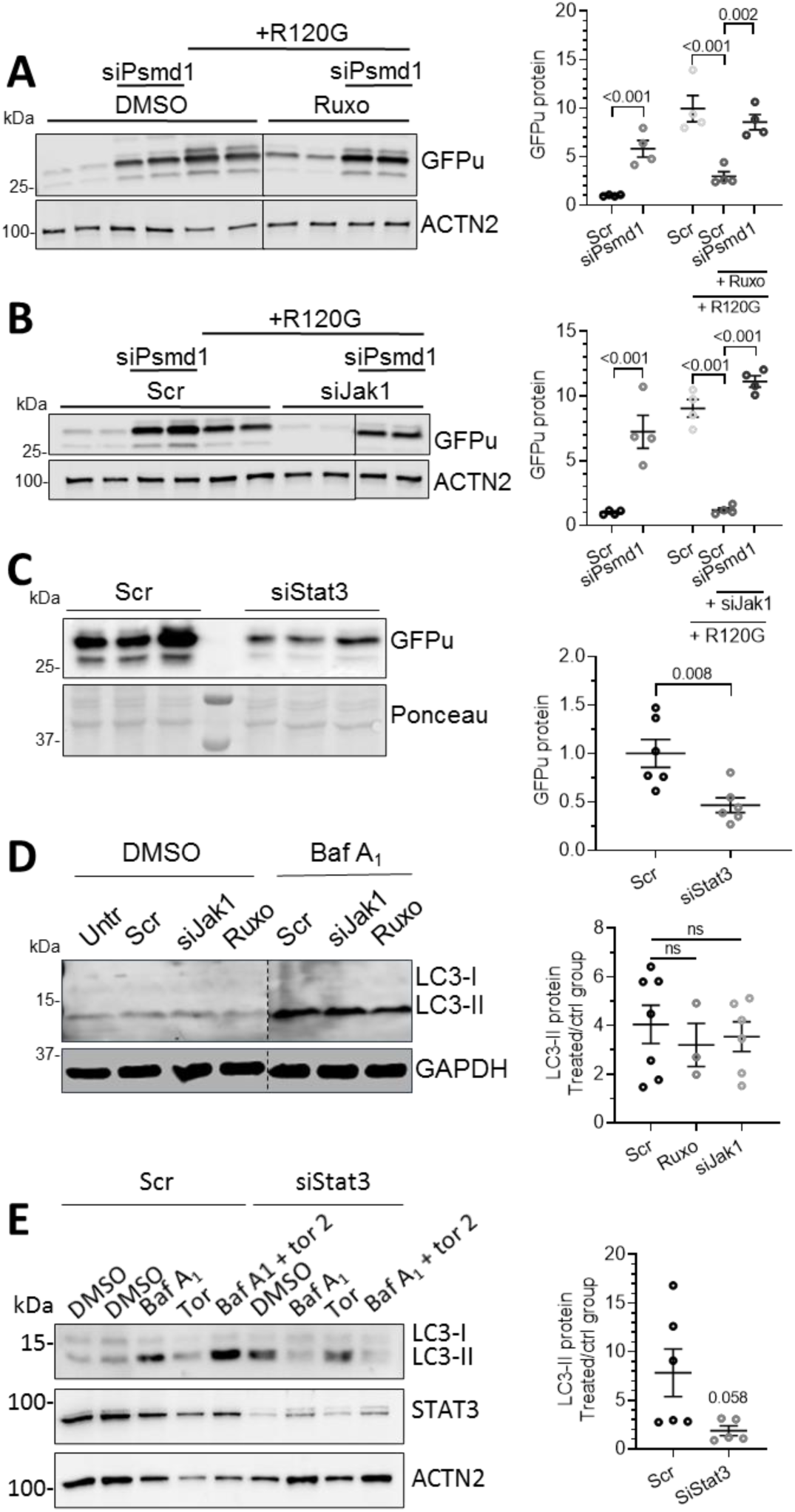
UPS-mediated degradation is enhanced by ruxolitinib treatment, *Jak1* or *Stat3* knockdown, but autophagic flux remains unaltered. NRVMs were transduced with AdV5-CMV-GFPu and/or AdV5-CMV-CRYAB^R120G^ for 5 days where indicated. **A**, NRVMs transfected with 20 nM siPsmd1 or scramble siRNA (scr) and treated with 3 µM ruxolitinib (ruxo) or DMSO for 5 days (every other day). **B**, NRVMs transfected with 20 nM siPsmd1 or scr and 100 nM siJak1 or scr. **C**, NRVMs transfected with 100 nM siStat3 or scr. **D,** NRVMs were transfected with 100 nM siJak1 or scr and/or treated with 3 µM ruxolitinib (ruxo). Before harvest, cells were treated with 50 nM bafilomycin A1 (baf A1) or DMSO for 2.5 h. **E**, NRVMs transfected with 100 nM siStat3 or scr. Before harvest, cells were treated with 50 nM bafilomycin A1 (baf A1) and/or torin 2 for 2.5 h. **A, B, C, D, E,** WBs were stained with antibodies against indicated proteins of protein extracts from treated NRVMs. WB quantification with Image Studio^TM^ or Image Lab^TM^ software. Data were obtained from 1 (A and B) or two (C-E) NRVM preparations with at least 3 wells per condition and are depicted as mean ± SEM, and p-values were obtained with the two-way ANOVA and Tukey’s multiple comparisons post-hoc or with the unpaired Student’s t-test (**C**). Abbreviation: ns, non-significant.

To assess ALP-mediated degradation, NRVMs were treated with the lysosomal V-ATPase inhibitor bafilomycin A_1_ or DMSO to assess the autophagic flux via the quantification of the autophagy marker LC3-II by Western blot analysis. As expected, bafilomycin A_1_ treatment resulted in a marked accumulation of LC3-II and a delta of ∼3 for the autophagic flux (Figure 5D). Ruxolitinib treatment slightly reduced (not significant) the autophagic flux (delta ∼1.8; Figure 5D). In line with our previous study,^12^ *Jak1* knockdown did not affect the autophagic flux (Figure 5D) or autophagy activation via mammalian target of rapamycin complex 1 (mTORC1) signaling (Figure S5). This experiment was also repeated with *Stat3* knockdown and the combination of bafilomycin A_1_ and torin 2, an autophagy activator to exaggerate the autophagic flux (Figure 5E). With *Stat3* knockdown the autophagic flux was reduced but not significantly (Figure 5E). These results suggest that CRYAB p.Arg120Gly aggregate removal with ruxolitnib treatment is independent of autophagic activity, but may be dependent on UPS activity.

### Knockdown of essential proteasomal subunit PSMD1 results in CRYAB p.Arg120Gly aggregate accumulation with ruxolitinib treatment or Jak1 knockdown

To evaluate if the ruxolitinib-enhanced UPS-mediated degradation is responsible for the lower CRYAB p.Arg120Gly aggregated load, NRVMs were transduced with AdV5-CRYAB^R120G^-GFP and transfected with siPsmd1 to block UPS-mediated degradation in the presence of ruxolitinib (Figure 6). *Psmd1* knockdown did not lead to further CRYAB p.Arg120Gly aggregate accumulation in DMSO-treated NRVMs (Figure 6A), probably due to the already impaired UPS by the aggregates. In contrast, in ruxolitinib-treated NRVMs, *Psmd1* knockdown resulted in CRYAB p.Arg120Gly aggregate accumulation (Figure 6A), indicating that the ruxolitinib-induced enhancement of UPS-mediated degradation is responsible for the removal of the CRYAB p.Arg120Gly aggregates. The experiment was repeated with *Jak1* knockdown with similar results (Figure 6B), supporting the findings.

**Figure 6.**
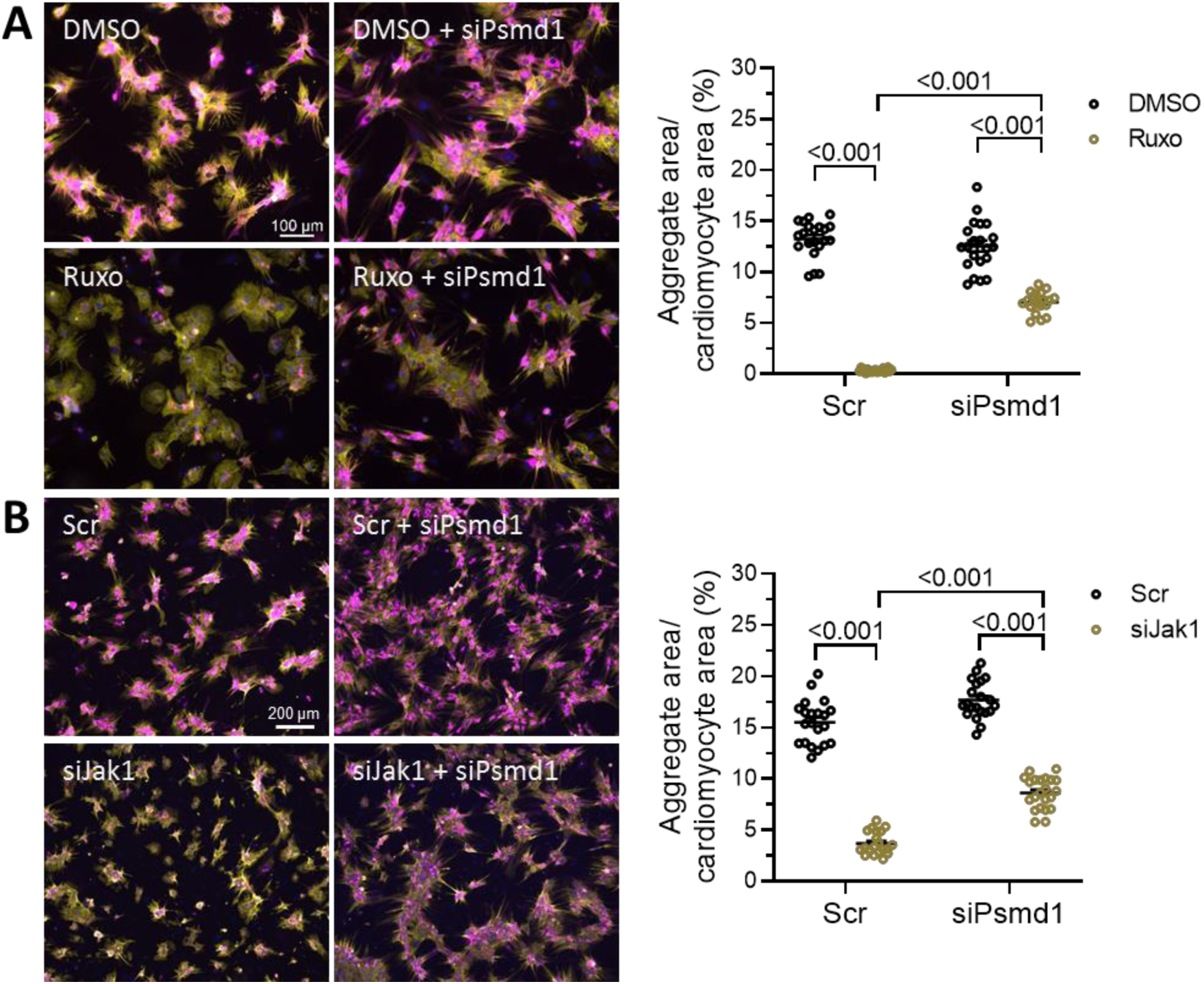
Knockdown of essential proteasomal subunit *Psmd1* results in CRYAB p.Arg120Gly aggregate accumulation with ruxolitinib treatment or *Jak1* knockdown. NRVMs transfected with 20 nM siPsmd1 or scramble siRNA (scr), transduced with Ad5-CMV-CRYAB^R120G^, treated with 3 µM ruxolitinib (ruxo) or DMSO (**A**), or transfected with 100 nM siJak1 or scr (**B**) and fixed after 6 days. **A**, **B**, Representative immunofluorescence images; Scale bar = 100 µm (A) or 200 µm (B). Aggregates are depicted in magenta (CRYAB^R120G^-GFP), cardiomyocytes in yellow (anti-cardiac troponin I) and nuclei in blue (DAPI). Quantification of aggregates in cardiomyocytes with NIS Elements software. Data were obtained from 3 independent NRVM preparations with at least 3 wells per condition and at least 5 images per well and are depicted as mean ± SEM, and the p-values were obtained with the two-way ANOVA and Tukey’s multiple comparisons post-hoc analysis.

### Jak1 knockdown regulates gene expression in NRVMs including gene expression of muscle-relevant E3 ubiquitin ligases

Since the JAK-STAT pathway regulates gene expression, paired-end RNAseq was performed on RNA extracts from NRVMs transfected with siRNA directed against *Jak1* or scramble siRNA. Data were analyzed with a differential gene expression analysis method. As a result, 117 RNAs were upregulated and 92 downregulated after *Jak1* knockdown (Figure S6A). As expected with modulation of the JAK-STAT pathway, expression of genes involved in biological processes such as cell differentiation, cytokine production, and inflammatory response (Figure S6B), molecular functions such as phosphotransferase activity, chemokine activity, and receptor binding (Figure S6C), and pathways such as interferon-gamma and interleukin signaling (Figure S6D) were primarily affected. When mapping the identified RNAs to data sets^21^ with gene identifiers of proteins involved in the UPS or ALP, a match with 5 ubiquitinating enzymes was found (Figure S6E). In addition, RNAseq results were analyzed for important muscle E3 ubiquitin ligases^23^ (Table 1) and we found significantly higher RNA levels for ankyrin repeat and SOCS box-containing 2 (*Asb2*), F-box protein 40 (*Fbxo40*), membrane-associated ring-CH-type finger 3 (*March3*), receptor-associated protein of the synapse (*Rapsn*), ring finger protein 152 (*Rnf152*), ring finger protein 207 (*Rnf207*), tripartite motif containing 50 (*Trim50*) and a tendency to higher RNA levels for F-box and leucine-rich repeat 21 (*Fbxl21*), and muscle-specific ring finger protein 2 (*Murf2*/*Trim55*), suggesting UPS activation through upregulating gene expression of E3 ubiquitinating ligases. Apart from that, NanoString RNA analysis of UPS, ALP and basic cardiomyocyte panels for ruxolitinib- or DMSO-treated NRVMs revealed higher RNA levels of the E3 ubiquitinating ligases *Murf1* (*Trim63*) and *Murf2* (*Trim55*) and lower RNA levels of the deubiquitinating enzyme ubiquitin carboxy-terminal hydrolase L1 (*Uchl1*) with ruxolitinib treatment (Figure S7A), and slightly lower RNA levels of lysosomal proteases cathepsin D (*Ctsd*) and cathepsin H (*Ctsh*; Figure S7B). In addition, gene expression of several cardiomyocyte-relevant proteins was altered such as four-and-half LIM domains protein 1 (*Fhl1*), α-myosin heavy chain (*Myh6*), natriuretic peptide A (*Nppa*), natriuretic peptide B (*Nppb*) and ryanodine receptor 2 (*Ryr2*; Figure S7C), indicating specific gene regulation of the JAK-STAT pathway in cardiomyocytes. Looking at JAK-STAT pathway members, expression levels of *Jak1* and *Stat1* were not affected (Figure S7D), and *Stat3* levels were about 50% lower with ruxolitinib treatment (Figure S7D).

**Table 1.**
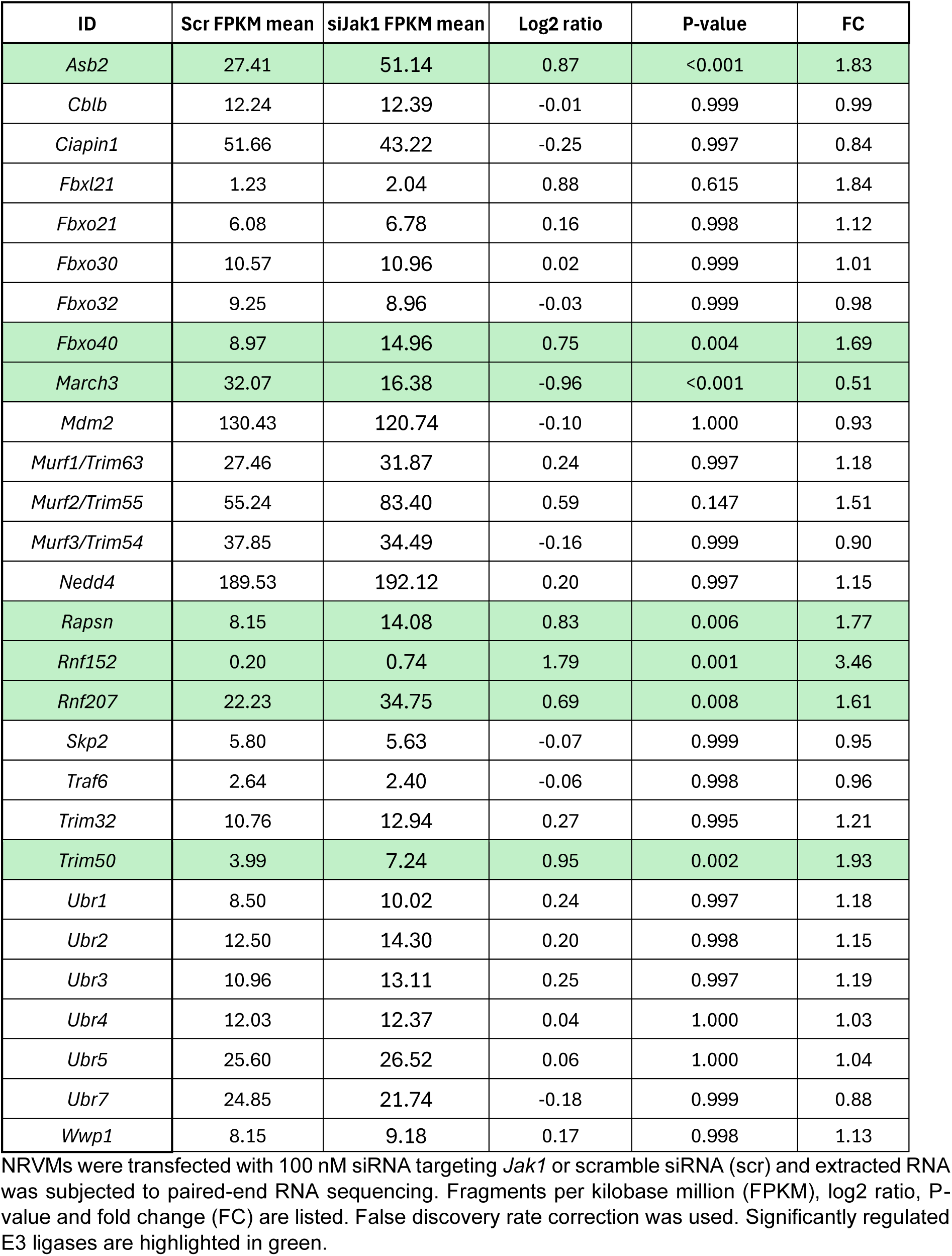
*Jak1* knockdown results in higher expression levels of important muscle E3 ligases.

### Knockdown of E3 ligase Asb2 abolishes the effect of ruxolitinib on CRYAB p.Arg120Gly aggregates

To test if the upregulation of the muscle-relevant E3 ligases was responsible for the clearance of the CRYAB p.Arg120Gly aggregates, we performed siRNA knockdown experiments of *Asb2*, *Rapsn*, *Trim50,* and *Rnf207* in combination with ruxolitinib. Rnf152, March3 and Fbxo40 were not tested due to low expression levels, downregulation by ruxolitinib, or failed siRNA knockdown, respectively. SiRNAs targeting *Asb2*, *Rapsn*, *Trim50,* and *Rnf207* demonstrated mRNA knockdown >50% (data not shown). Whereas *Raspn*, *Trim50,* and *Rnf207* knockdown did not reveal any difference in CRYAB p.Arg120Gly aggregate load after ruxolitinib treatment (Figure S8), *Asb2* knockdown abolished the effect of ruxolitinib on CRYAB p.Arg120Gly aggregates completely (Figure 7 A and B). Since Asb2β targets desmin for proteasomal degradation,^24^ we tested ruxolitinib and *Jak1* siRNA on desmin levels in NRVMs and found that they were 2-fold lower (Figure S9).

**Figure 7.**
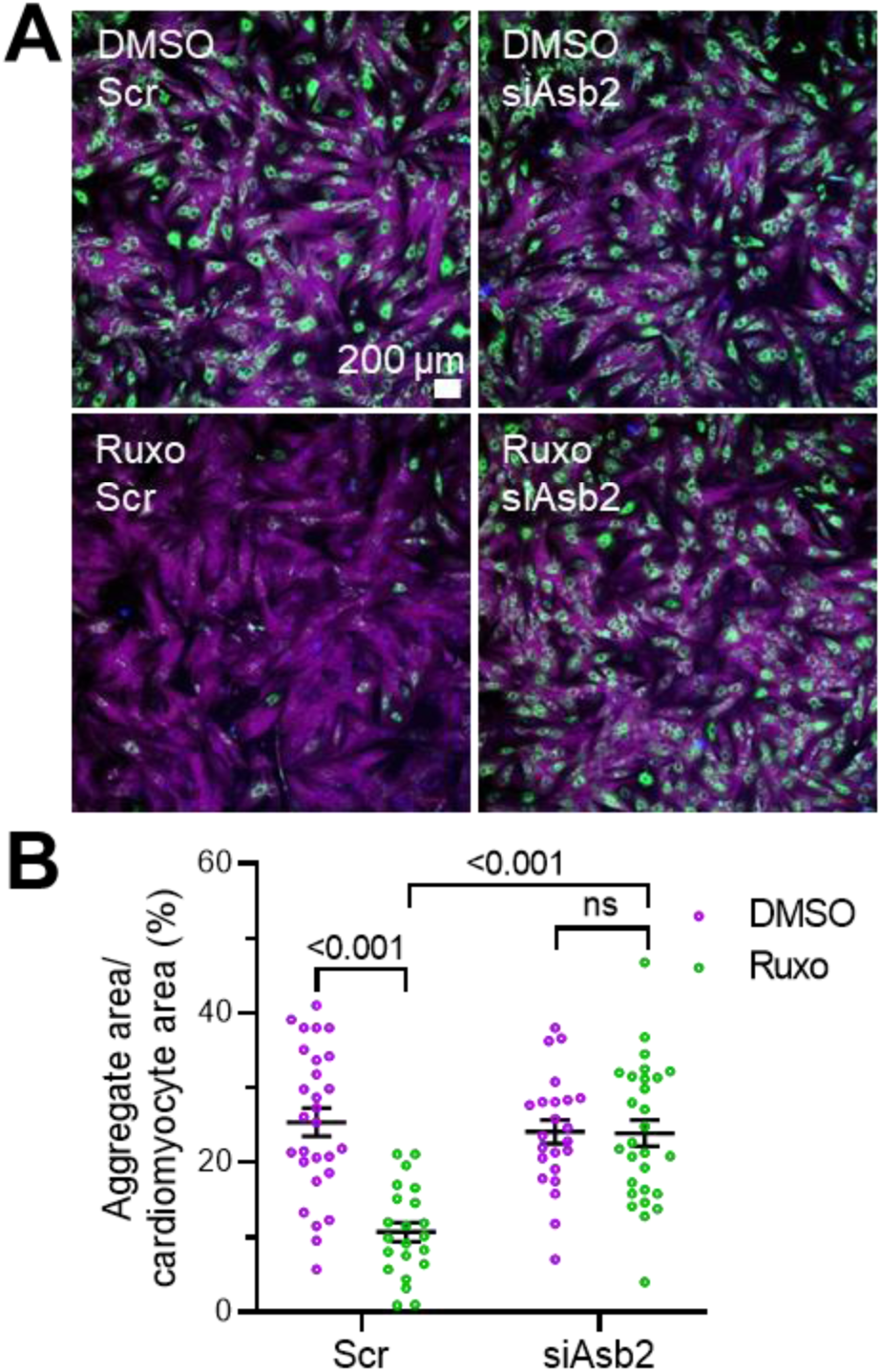
*Asb2* knockdown abolishes ruxolitinib effect on CRYAB p.Arg120Gly aggregates. **A**, NRVMs transfected with 50 nM siAsb2 or scramble siRNA (scr), transduced with Ad5-CMV-CRYAB^R120G^, treated with 3 µM ruxolitinib (ruxo) or DMSO and fixed after 5 days. Representative immunofluorescence images. Scale bar = 200 µm. Aggregates are depicted in green (CRYAB^R120G^-GFP), cells in purple (phalloidin) and nuclei in blue (DAPI). **B**, Quantification of aggregates in cells with NIS Elements software. Data were obtained from one NRVM preparation with at least 3 wells per condition and at least 6 images per well and are depicted as mean ± SEM, and p-values were obtained with the two-way ANOVA and Tukey’s multiple comparisons post-hoc analysis. Dots represent number of analyzed images. Abbreviation: ns, non-significant.

### Higher phosphorylated STAT3 protein levels in DRM mice with hypertrophy and dysfunction

To test if JAK-STAT signaling is pathologically altered in DRM mice, we analyzed the hearts of DRM mice and compared them to their non-transgenic (NTG) littermates. DRM mice did not exhibit cardiac hypertrophy at 1 month, whereas hypertrophy was fully established at 4 months and did not increase further with age (Figure 8A). Ejection fraction (EF) was lower in DRM mice at 7 but not 4 months of age (Figure 8B). Therefore, we determined phosphorylated STAT3 (P-STAT3) protein levels in 1-, 4- and 7-month-old mice. The P- STAT3/STAT3 ratio did not differ in 1-month-old, but was 2- and 5-fold higher in 4- and 7- month-old DRM than NTG mice, respectively (Figure 8C and D), suggesting pathological activation of the pathway with age. *Stat3* mRNA levels did not differ between the groups at 7 months (Figure 8E).

**Figure 8.**
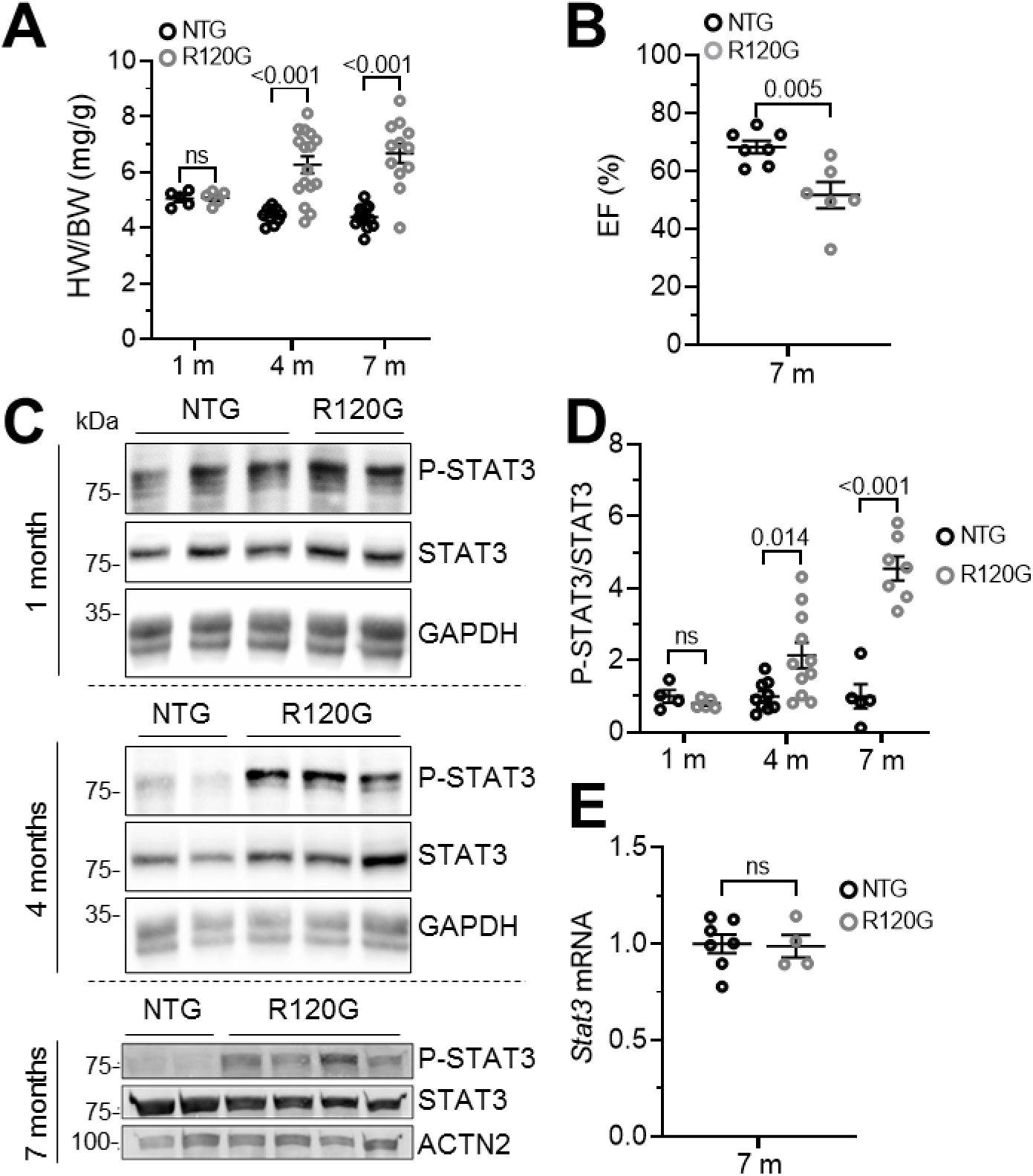
Phosphorylated STAT3 protein levels are higher in CRYAB p.Arg120Gly transgenic mice with hypertrophy. **A**, Heart weight-to-body weight ratio (HW/BW) in 1-, 4- and 7-month-old CRYAB p.Arg120Gly transgenic (R120G) and non-transgenic (NTG) mice. **B**, Ejection fraction (EF) in 7-month-old R120G and NTG mice. **C**, Representative Western blots of phosphorylated (P-)STAT3, STAT3 and indicated control in 1-, 4- and 7-month-old R120G and NTG mice. **D**, P-STAT3/STAT3 quantification of Western blots from 1-, 4- and 7-month-old R120G and NTG mice. **E**, *Stat3* mRNA level determined by RT-qPCR from 7-month-old R120G and NTG mice. Western blot quantification was performed with Image Lab software. Data are depicted as mean ± SEM, and p-values were obtained with the unpaired Student’s t-test. Abbreviation: ns, non-significant.

### Ruxolitinib treatment prevents the development of cardiac dysfunction in CRYAB p.Arg120Gly transgenic mice

To test if ruxolitinib blunts cardiac disease in DRM mice, 21-week-old DRM and NTG mice were treated twice daily with 75 mg/kg ruxolitinib or vehicle by oral gavage for 3 weeks with initial and final echo (Figure 9A). P-STAT3/STAT3 ratio was normalized by ruxolitinib treatment (Figure 9B). At the beginning of the experiment, cardiac function, represented by EF did not differ between DRM and NTG mice (Figure 9C). As anticipated, EF was lower in vehicle-treated DRM than NTG mice at the end of the experiment. Ruxolitinib treatment prevented the drop in EF in DRM mice (Figure 9C). Cardiac hypertrophy represented by LVM/BW and HW/BW was higher in DRM than NTG mice and not affected by ruxolitinib treatment (Figure 9C and D). Body weights did not differ between the groups (Figure 9D). CRYAB p.Arg120Gly aggregate load was lower in ruxolitinib than in control-treated mice (Figure 9E).

**Figure 9.**
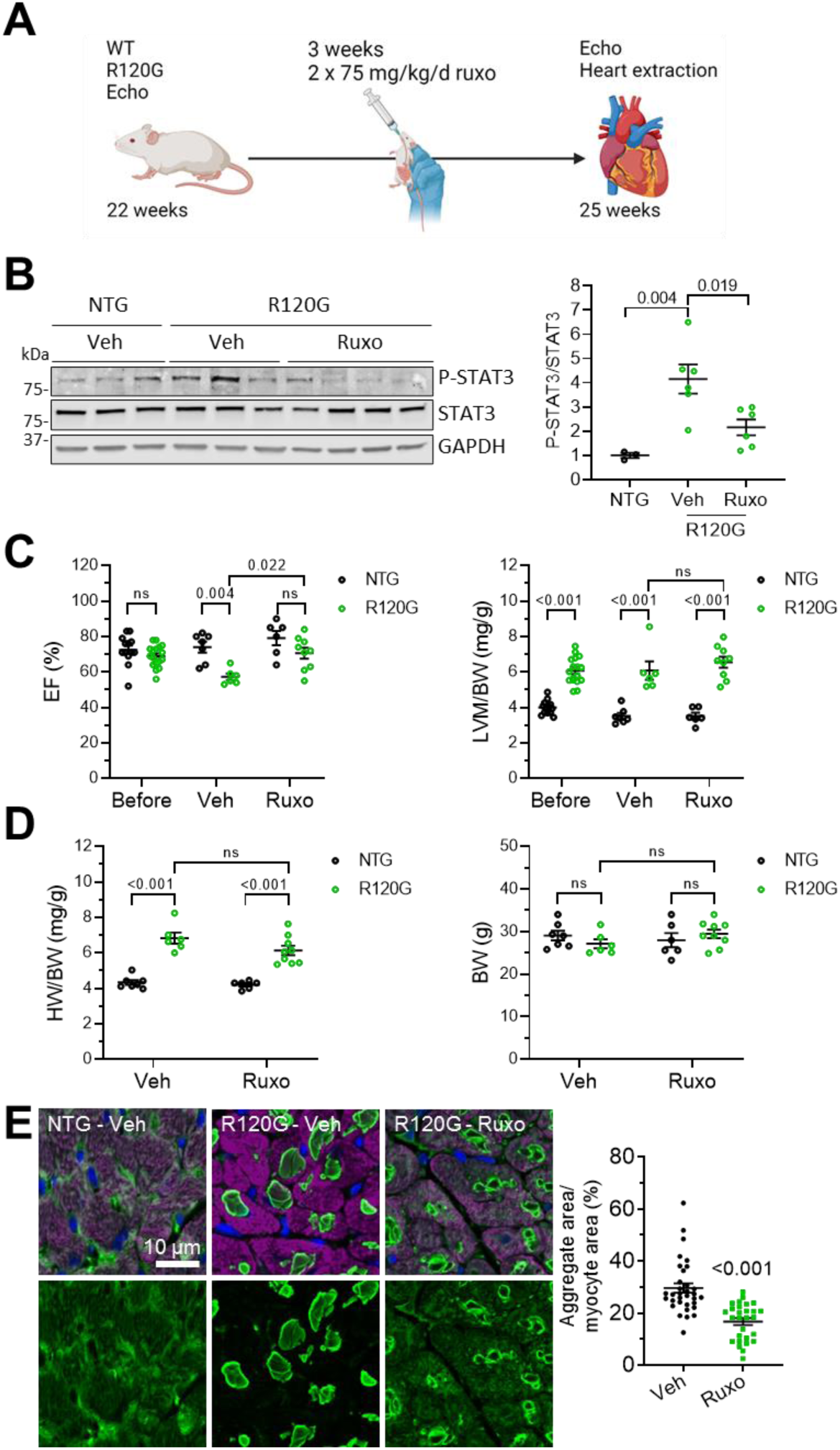
Ruxolitinib treatment of CRYAB p.Arg120Gly transgenic mice prevents development of cardiac dysfunction. CRYAB p.Arg120Gly transgenic (R120G) and non-transgenic (NTG) mice treated for 3 weeks with 75 mg/kg ruxolitinib (ruxo) or vehicle (veh) twice-daily oral gavage. Transthoracic echocardiography was performed at the start (before, 21-week-old) and the end of the treatment (veh or ruxo, 24-week-old). **A**, Scheme of experimental outline. **B**, WB and quantification of phosphorylated (P-) STAT3, STAT3 and indicated controls at the end of treatment. WB quantification was performed with Image Lab software. **C,** Ejection fraction (EF) and left ventricular mass-to-body weight ratio (LVM/BW) before and after (veh/ruxo) treatment. **D**, Heart weight-to-body weight ratio (HW/BW) and body weight (BW) at the end of the treatment. **E**, Representative images and quantification of R120G and NTG mouse heart sections after vehicle or ruxolitinib treatment. CRYAB is depicted in green, cardiomyocytes in purple (anti-cardiac troponin I) and nuclei in blue (DAPI). Scale bar = 10 µm. Quantification of aggregates in cardiomyocytes with NIS Elements software. At least 5 images of 3 mice per group were analyzed. Data are depicted as mean ± SEM, and p-values were obtained with the one-way (**B**) or two-way ANOVA and Tukey’s multiple comparisons post-hoc analysis (**C, D**) or unpaired Student’s t-test (**E**). Abbreviation: ns, non-significant.

### Conditional Jak1 KO prevents the development of cardiac dysfunction in CRYAB p.Arg120Gly transgenic mice

To test if *Jak1* KO blunts cardiac disease in DRM mice, 8-week-old DRM and NTG mice crossed with conditional *Jak1* KO and cardiomyocyte-specific αMHC-MerCreMer mice were fed with tamoxifen chow to induce KO and analyzed by echocardiography and heart extraction at 26 weeks of age (Figure 10A). *Jak1* mRNA levels were about 40% and 70% lower in induced heterozygous and homozygous *Jak1* KO mice, respectively (Figure 10B). JAK1 protein levels were about 70% lower in induced homozygous *Jak1* KO mice (Figure 10C). It should be noted that *Jak1* KO is only induced in cardiomyocytes and other cells of the heart still express *Jak1*. As anticipated, EF was lower in 26-week-old DRM than NTG mice with *Jak1* wt (Figure 10D). Strikingly, in the heterozygous and homozygous *Jak1* KO groups, EF did not differ between DRM and NTG mice (Figure 10D). EF did not differ between *Jak1* KO and wt in the NTG group, whereas it was higher in homozygous *Jak1* KO than in *Jak1* wt in DRM mice (Figure 10D). Cardiac hypertrophy represented by LVM/BW and HW/BW was higher in DRM than NTG mice with *Jak1* wt and not affected by *Jak1* KO (Figure 10D and E). Body weights did not differ between the groups (Figure 10E). CRYAB p.Arg120Gly aggregate load was lower in *Jak1* KO than in Jak1 wt mouse ventricular tissues (Figure 10F).

**Figure 10.**
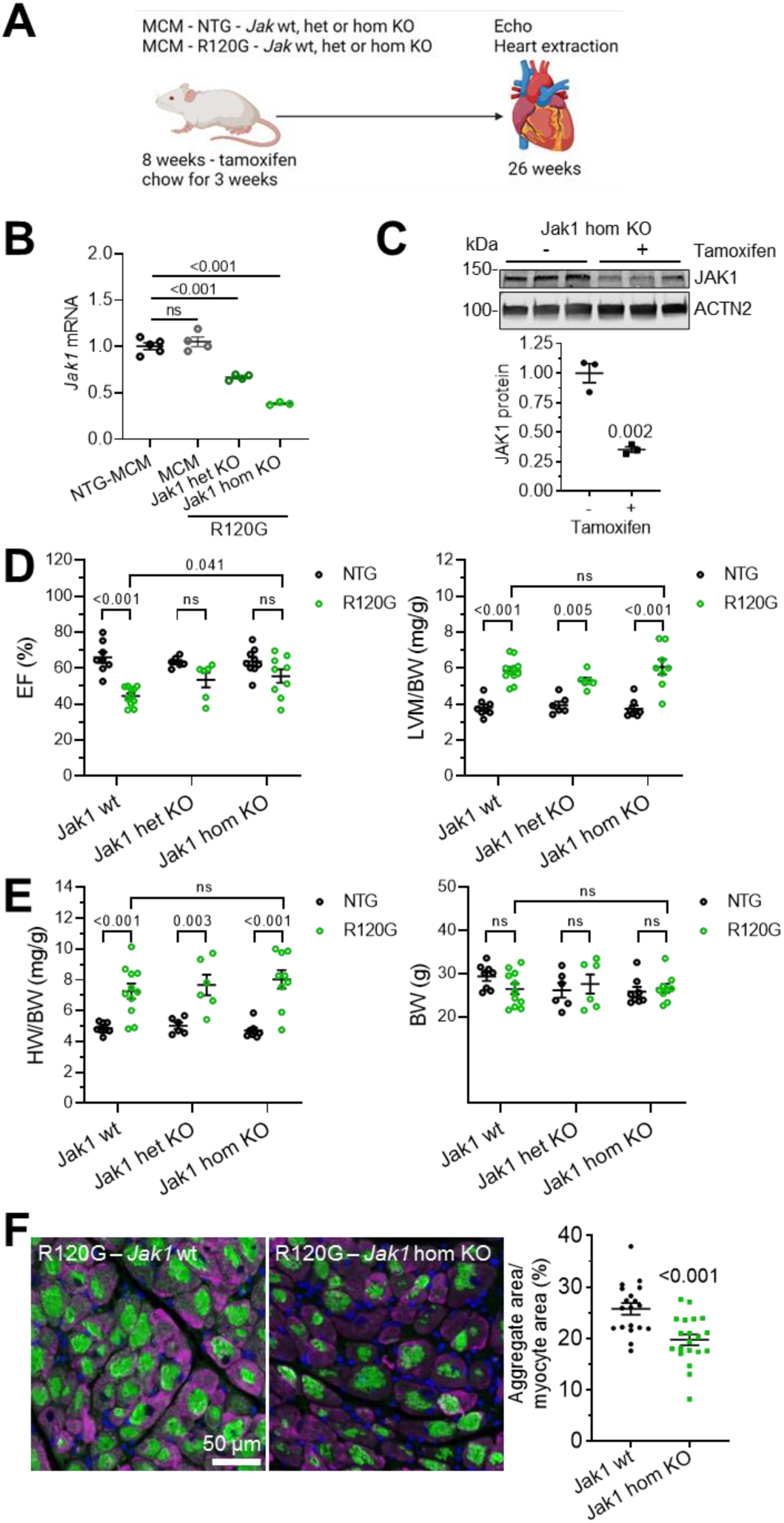
*Jak1* knockout prevents development of cardiac dysfunction in CRYAB p.Arg120Gly transgenic mice. Heterozygous (het) or homozygous (hom) *Jak1* knockout (KO) was induced in MCM-transgenic (TG) mice crossed with CRYAB p.Arg120Gly TG (R120G) or non-transgenic (NTG) mice with tamoxifen chow. Transthoracic echocardiography was performed 26 weeks and hearts were extracted. **A**, Scheme of experimental outline. **B**, *Jak1* mRNA levels determined by RT-qPCR. **C**, JAK1 protein levels determined by Western blot and normalized to ACTN2. Western blot quantification was performed with Image Lab software **D,** Ejection fraction (EF) and left ventricular mass-to-body weight ratio (LVM/BW). **E**, Heart weight-to-body weight ratio (HW/BW) and body weight (BW). **F**, Representative images and quantification of mouse heart sections. Aggregates are depicted in green, cardiomyocytes in purple (anti-cardiac troponin I) and nuclei in blue (DAPI). Scale bar = 50 µm. Quantification of aggregates in cardiomyocytes with NIS Elements software. At least 5 images of 3 mice per group were analyzed. Data are depicted as mean ± SEM, and p-values were obtained with the one-way (**B**) or two-way ANOVA (**D**, **E**) with Tukey’s multiple comparisons post-hoc analysis or unpaired Student’s t-test (**C**, **F**). Abbreviation: ns, non-significant.

## Discussion

In the present study, we tested the impact of JAK1 knockdown and inhibition in CRYAB p.Arg120Gly DRM models. Our main findings were: i) ruxolitinib treatment prevented the formation of and cleared CRYAB p.Arg120Gly aggregates in NRVMs and hiPSC-CMs; ii) knockdown of *Jak1* and *Stat3*, but not *Jak2* prevented the formation of CRYAB p.Arg120Gly aggregates; iii) ruxolitinib treatment resulted in higher UPS function and higher RNA levels of muscle-relevant E3 ubiquitin ligases; iv) blocking UPS function blunted the effect of ruxolitinib on CRYAB p.Arg120Gly aggregates and v) JAK1 loss-of-function prevented cardiac dysfunction in DRM mice.

Ruxolitinib, first FDA-approved in 2011, is used to treat myelofibrosis, polycythemia vera, and graft versus host reaction in allogeneic stem cell transplantation. Furthermore, repurposing of ruxolitinib is tested for the treatment of COVID and autoimmune diseases.^25, 26^ Ruxolitinib competitively inhibits the ATP-binding site of JAK1 and JAK2 (IC_50_ = 3.3 nM and 2.8 nM in cell-free assay, respectively).^27^ Reported adverse events with ruxolitinib include peripheral edema, cardiovascular events, diarrhea, infections, and hematologic/inhibition of hematopoiesis, and a regular check of blood count and kidney function is necessary.^28, 29^ In the past years, there has been a race to identify JAK inhibitors which are more specific to one member of the family, though with limited success. In this study, we tested 3 inhibitors, solcitinib (not approved), upadacitinib (FDA-approved), and filgotinib (FDA-approved), which were primarily tested for inhibiting mutant JAK1 in diseases such as rheumatoid arthritis or inflammatory bowel disease. We found that all of them efficiently reduced CRYAB p.Arg120Gly aggregate load in NRVMs and hiPSC-CMs. Since none of the available inhibitors is entirely specific to JAK1, we performed siRNA knockdown experiments and found that *Jak1* but not *Jak2* knockdown prevented CRYAB p.Arg120Gly aggregate accumulation, indicating distinct functional differences of JAK1 and JAK2 in cardiomyocytes. Furthermore, *Jak1* and *Stat3* mRNA levels were both markedly higher than the other JAK and STAT members in NRVMs and hiPSC-CMs, suggesting an important role for them in cardiomyocytes. Especially, *Jak3*, *Stat4*, *Stat5a,* and *Stat5b* mRNA levels were very low or not detected in NRVMs and hiPSC-CMs and may therefore be less important than the other JAK-STAT family members in cardiomyocytes. In line with this, whole-body knockouts of *Jak1*, *Jak2,* and *Stat3* are lethal in mice, whereas knockouts of the other JAK-STAT pathway members are viable with defects in immune response and/or hematopoiesis.^30^

Variants in JAKs can cause severe diseases such as rheumatoid arthritis and inflammatory bowel diseases for *JAK1* variants, myeloproliferative neoplasms for *JAK2* variants, and severe immune deficiency for *JAK3* and *TYK2* variants.^14^ Similarly, several *STAT* variants increase the incidence of malignancies, infections, and autoimmune diseases.^31^ The role of JAK1 in cardiac health and disease remains relatively obscure. For STAT3, several studies pointed out its importance in the heart. For example, cardioprotective roles have been described such as in cardiomyocyte-specific *Stat3* knock-out in mice, which were more susceptible to ischemia/reperfusion injury, had reduced cardiac function, and increased mortality after myocardial infarction and with age.^32, 33^ On the other hand, STAT3 has been described to stimulate cardiac hypertrophy in experimental models.^34, 35^ However, the role of STAT3 in cardiomyocyte hypertrophy was questioned when interleukin-6 deletion, reducing STAT3 signaling did not affect the development of hypertrophy.^36^ In human hearts, higher^37^ or lower^38^ P-STAT3 levels was observed in dilated cardiomyopathy. In this study, we found that the steady-state level of P-STAT3 was higher in DRM mice.

Apart from the canonical JAK-STAT signaling, a few non-canonical functions have been proposed for JAK1. For example, direct epigenetic regulation of gene expression by H3 phosphorylation through JAK has been described.^39^ Furthermore, one study showed that JAK1 regulates many genes in response to ER stress and coimmunoprecipitates with ATF4.^40^ Yet another study found that ER stress stimulated JAK1-dependent STAT3 activation.^41^ Here, we provide evidence for a novel non-canonical function of JAK1-STAT3 in the regulation of the UPS. The UPS is responsible for the specific physiological and pathological degradation of many proteins including desmin and CRYAB.^42^ For a long time, it has been assumed that the UPS can only remove single proteins, but not larger protein aggregates. However, it has been shown that protein aggregates are removed by the UPS in the nucleus, where there is no access to the autophagic machinery,^43^ and in the cytosol of autophagy-defective *Atg5* knockout cells.^44^ Here, we show that ruxolitinib treatment or *Jak1* knockdown reduces CRYAB p.Arg120Gly aggregates, enhances UPS function, leads to higher expression of E3 ubiquitin ligases including *Asb2*, and does not enhance autophagic activity or alter expression of autophagy-related genes, suggesting that the CRYAB p.Arg120Gly aggregates are cleared by the UPS. Several studies have shown that the upregulation of UPS or ALP function can prevent the formation of CRYAB p.Arg120Gly aggregates,^10, 45, 46^ but did not show the clearance of pre-existing aggregates. This is important because the aggregates are already present when patients develop symptoms and seek health care. Important of note, our experiments suggest that ruxolitinib can clear small and medium-sized pre-existing CRYAB p.Arg120Gly aggregates. Therefore, a potential treatment would have to start relatively early in the disease.

Ruxolitinib treatment or *Jak1* KO in DRM mice underlined its therapeutic potential in vivo, since both led to lower CRYAB p.Arg120Gly aggregates and prevented a drop in cardiac function. Cardiac hypertrophy was not affected, which is in line with the finding that hypertrophy does not differ between 4- and 7-month-old DRM mice although the aggregate load increases. There may be a certain threshold of aggregate load to establish hypertrophy. In conclusion, it will be of great interest to further test if repurposing of JAK1 inhibitors is a potential therapy for DRM caused by CRYAB p.Arg120Gly. Also, testing the effect of ruxolitinib or other more specific JAK1 inhibitors not only in DRM, but in other protein aggregate-driven diseases could be of benefit. Furthermore, the novel pathway of ruxolitinib regulating the UPS will be of interest to all areas in which it is currently used and pave the way for other yet unconsidered applications.

## Acknowledgements

This work would have not been possible without and is dedicated to Jeffrey Robbins (CCHMC, Cincinnati, Ohio, USA), an outstanding scientist and mentor, who tragically passed away in 2022.

We thank Birgit Klampe, Thomas Schulze, Thomas Eschenhagen and Arne Hansen (UKE, Hamburg, Germany) for help with and providing hiPSC-CMs, Kritton Shay-Winkler (CCHMC, Cincinnati, Ohio, USA) for preparation of NRVMs, the UKE HEXT Vector facility (Ingke Braren) for virus productions and help with live cell imaging, UKE NanoString core Facility for RNA analysis (Elisabeth Krämer, Saskia Schlossarek), Niels Pietsch (UKE, Hamburg, Germany) for help with dot plots and the CCHMC DNA Sequencing and Genotyping Core for RNA sequencing and analysis.

## Abbreviations

AdV5: Adenovirus serotype 5
ALP: Autophagy-lysosomal pathway
CRYAB: αB-crystallin
DRM: Desmin-related cardiomyopathy
HiPSC-CMs: Human-induced pluripotent stem cell-derived cardiomyocytes
NRVM: Neonatal rat ventricular myocytes
JAK: Janus kinase
STAT: Signal transducer and activator of transcription
SiRNA: Small interfering RNA
UPS: Ubiquitin-proteasome system

