## Supplemental figures for "Ruxolitinib clears CRYAB p.Arg120Gly aggregates through the ubiquitin-proteasome system"

**Table S1. RNA counts from RNA sequencing of JAK-STAT family members in NRVMs and hiPSC-CMs.**

| RNA ID | NRVMs FPKM | HiPSC-CMs RPKM |
|---|---|---|
| *Jak1/JAK1* | 37 | 3713 |
| *Jak2/JAK2* | 15 | 610 |
| *Jak3/JAK3* | 9 | 23 |
| *Tyk2/TYK2* | 5 | 1778 |
| *Stat1/STAT1* | 26 | 4009 |
| *Stat2/STAT2* | 17 | 3025 |
| *Stat3/STAT3* | 72 | 5452 |
| *Stat4/STAT4* | 0 | 210 |
| *Stat5a/STAT5A* | 2 | 112 |
| *Stat5b/STAT5B* | 7 | 854 |
| *Stat6/STAT6* | 14 | 1830 |
| *Mybpc3/MYBPC3* | 254 | 41649 |
| *Ryr2/RYR2* | 25 | 5822 |
| *Akt1/AKT1* | 93 | 4016 |
| *Mapk1/MAPK1* | 49 | 5289 |
| *Mapk3/MAPK3* | 45 | 531 |

Mean RNA counts of *Mybpc3*, *Ryr2*, *Akt1*, *Mapk1* and *Mapk3* are depicted as expression reference values (FPKM – fragments per kilobase million; RPKM – reads per kilobase million). High and low expression values within a family are depicted in blue and red shades, respectively.

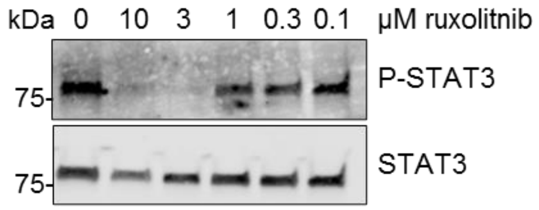


**Figure S1. Ruxolitinib concentration-response-curve on phosphorylated STAT3 levels in NRVMs.** NRVMs were treated with indicated concentrations of ruxolitinib or 0.1% DMSO. Western blot of protein extracts from treated NRVMs was stained with antibodies directed against phosphorylated (P-) STAT3 and STAT3.

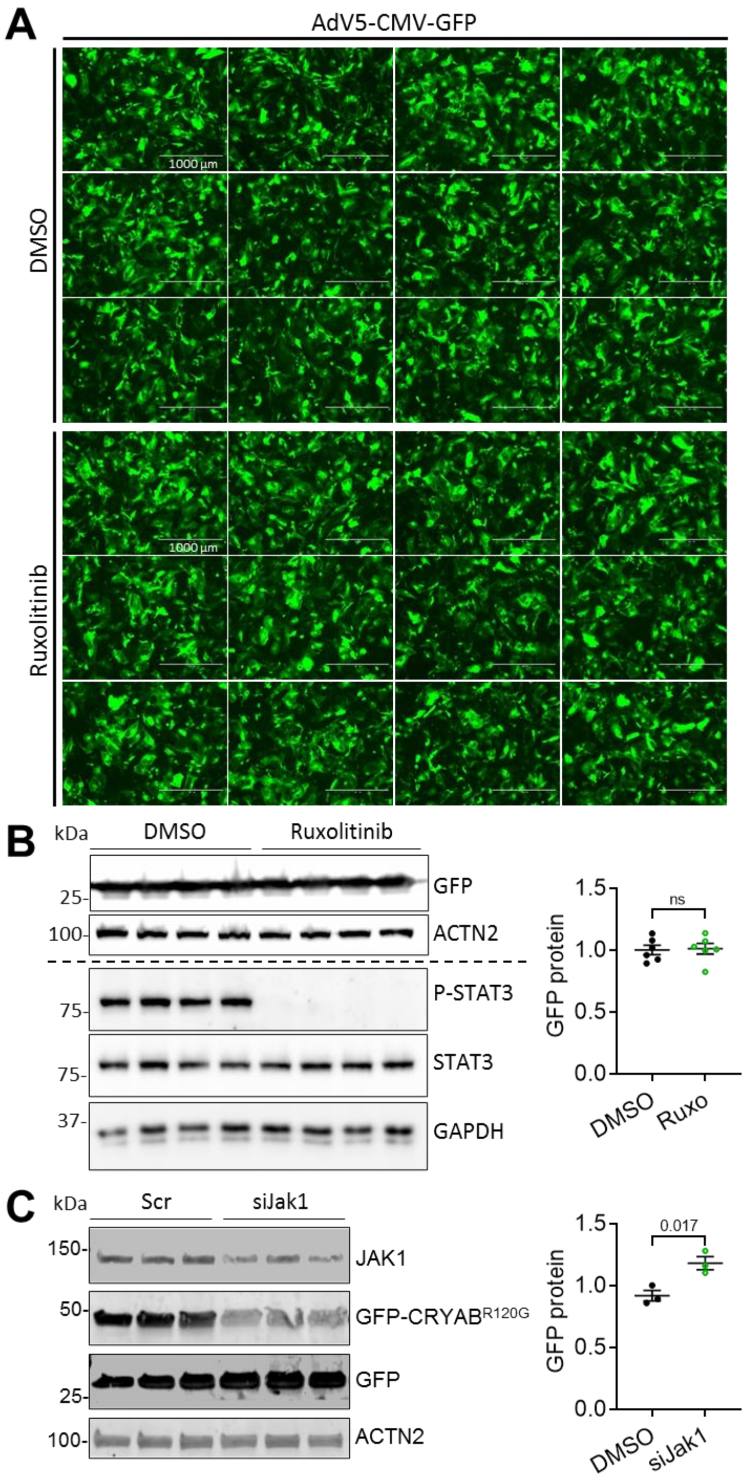


**Figure S2. Ruxolitinib or siJak1 treatment does not result in lower GFP levels in immunofluorescence or -blot.** NRVMs were transduced with AdV5-CMV-GFP (A-C) and AdV5-CMV-CRYAB$^{R120G}$-GFP (C) and treated with either 3 µM ruxolitinib or DMSO directly after transduction (A, B) or transfected with siRNA targeting JAK (siJak1) or scramble siRNA (scr) directly before transduction (C). Medium change with ruxolitinib or DMSO was performed every other day and cells were fixed or harvested 5 days after transduction. **A**, IF images. Scale bar = 1000 µm. **B**, **C**, Western blots of protein extracts from treated NRVM were stained with antibodies directed against indicated proteins. Quantification was performed with Image Lab (B) or Image Studio (C) software. Data are depicted as mean ± SEM, and p-value was obtained with the unpaired Student’s t-test. Abbreviation: ns, non-significant.

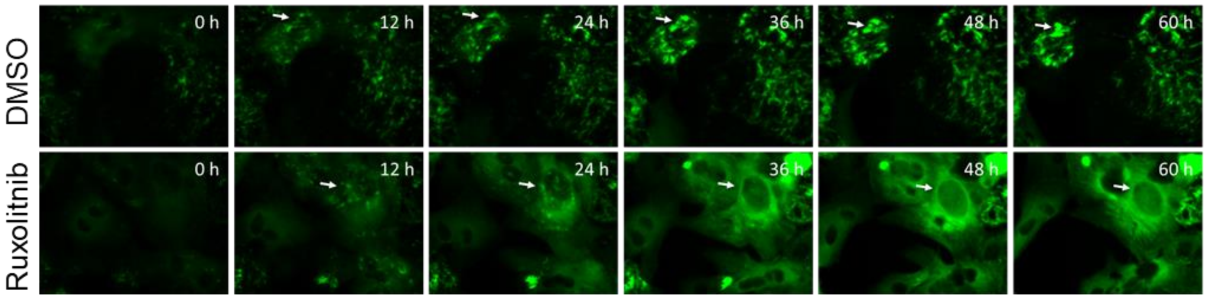


**Figure S3. Ruxolitinib dissolves CRYAB p.Arg120Gly aggregates in hiPSC-CMs.** Live cell imaging of hiPSC-CMs transduced with Ad5-CMV-CRYAB$^{R120G}$. Representative fluorescence images of videos from hiPSC-CMs treated with 3 µM ruxolitinib or 0.1% DMSO taken with a Nikon Biostation IM-Q time lapse imaging system (widefield). Aggregates are depicted in green (CRYAB$^{R120G}$-GFP). Arrows mark growing or dissolving aggregates. Corresponding videos can be found in online supplements.

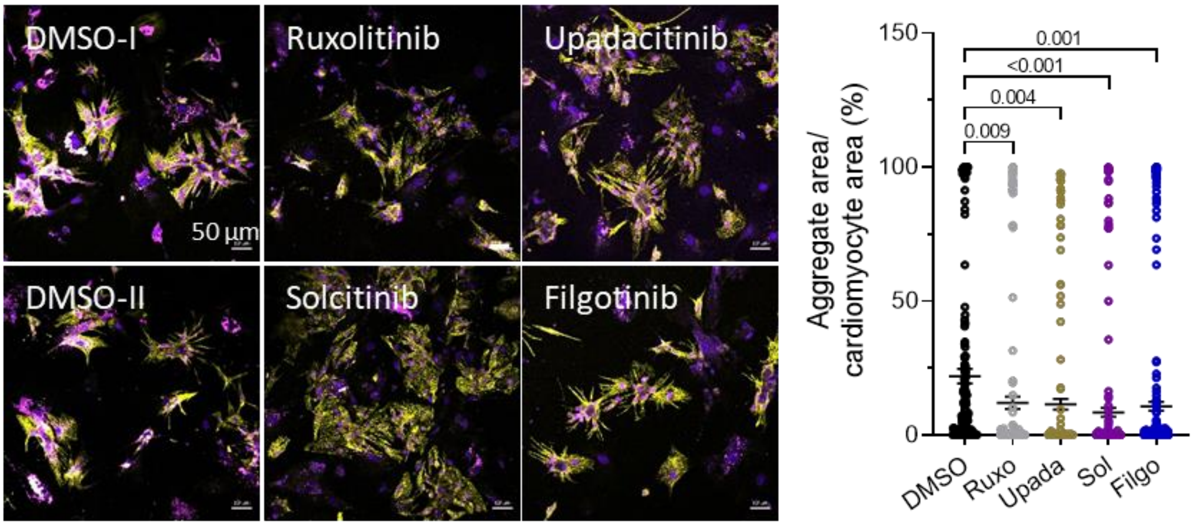


**Figure S4. Treatment with JAK1 inhibitor upadacitinib, solcitinib or filgotinib results in markedly lower CRYAB p.Arg120Gly aggregate load in NRVMs.** Representative immunofluorescence images of NRVMs treated with 3 µM ruxolitinib or 1 µM upadacitinib, solcitinib or filgotinib or 0.1% DMSO, transduced with AdV5-CMV-CRYAB$^{R120G}$-GFP and fixed after 4-6 days. Aggregates are depicted in magenta (CRYAB$^{R120G}$-GFP), cardiomyocytes in yellow (anti-cTnI), and nuclei in blue (DAPI). Quantification of aggregates in cardiomyocytes (NRVMs) with ImageJ software. Scale bar = 50 µm. Data were obtained from 2 independent NRVM preparations with at least 4 wells per condition and 5-8 images per well and are depicted as mean ± SEM, with p-values obtained with the one-way ANOVA and Dunnett's post-hoc analysis.

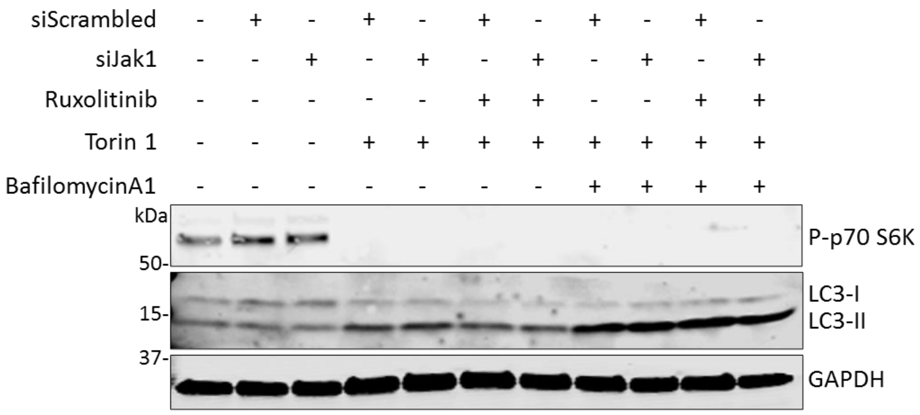


**Figure S5.** ***Jak1*** **knockdown or ruxolitinib treatment has no effect on mTORC1 signaling or LC3 levels in NRVMs.** NRVMs were transfected with siRNA targeting *Jak1* or scrambled siRNA and treated as indicated with ruxolitinib, torin 1 (autophagy activator) and/or bafilomycin A1 (autophagy inhibitor). Western blot of protein extracts from treated NRVMs was stained with antibodies directed against phosphorylated (P-) p70 S6K (mTORC1 target), LC3 (autophagy marker) and GAPDH (loading control).

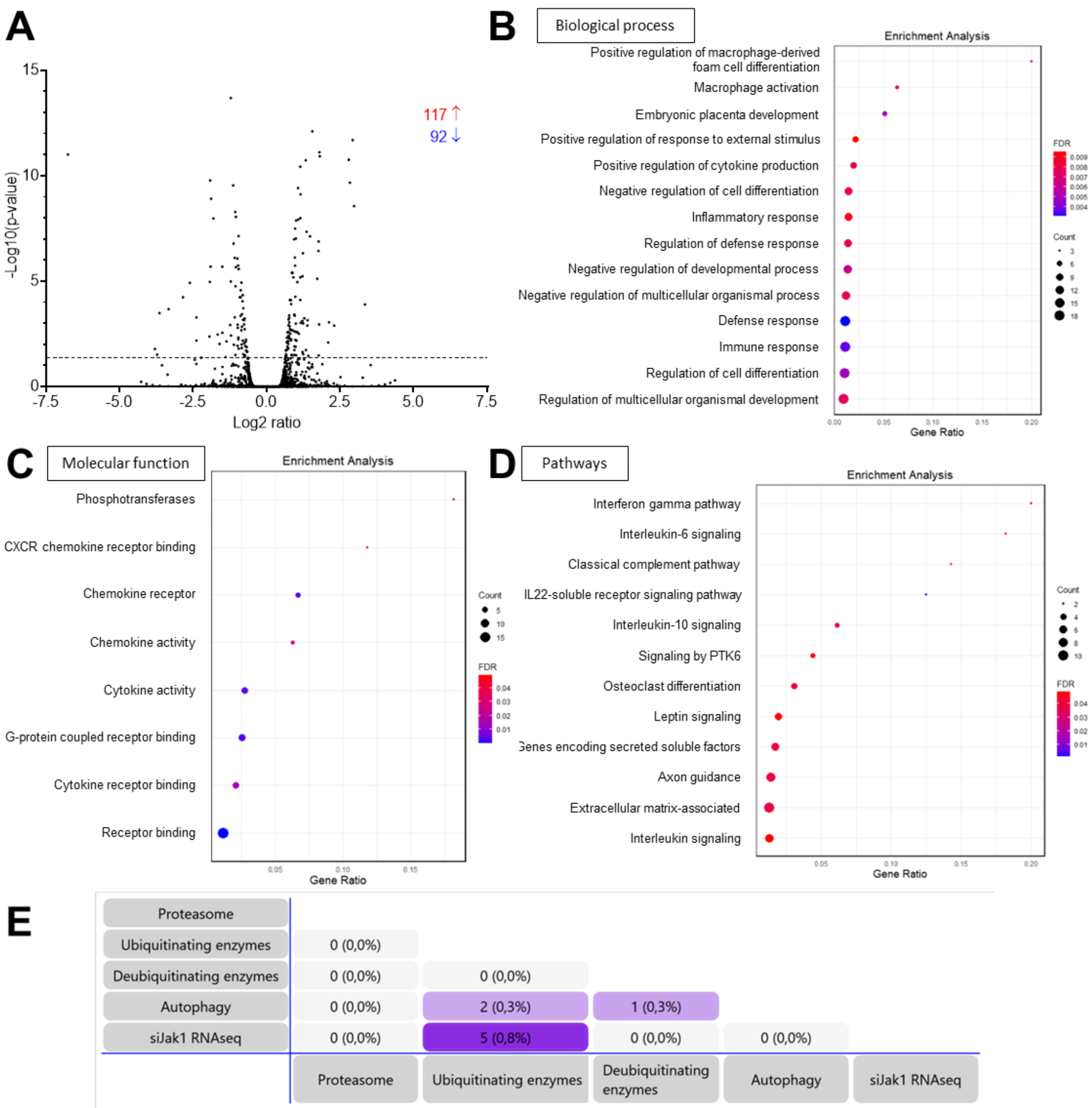


**Figure S6. Effects of *Jak1* knockdown on RNA levels in NRVMs.** NRVMs were transfected with 100 nM siRNA targeting *Jak1* or scramble siRNA and extracted RNA was subjected to paired-end RNA sequencing. Data were obtained from 3 samples of 3 independent NRVM preparations. **A**, Volcano plots show the -log10 of P-value vs. the magnitude of change (log2 ratio) of mRNA levels in siJak1/scr. Dot plots of enrichment analysis of **B**, biological processes, **C**, molecular function, and **D**, pathways. **E**, Mapping of significantly up- or- downregulated RNAs to gene identifiers related to proteasome, ubiquitinating enzymes, deubiquitinating enzymes and autophagy.

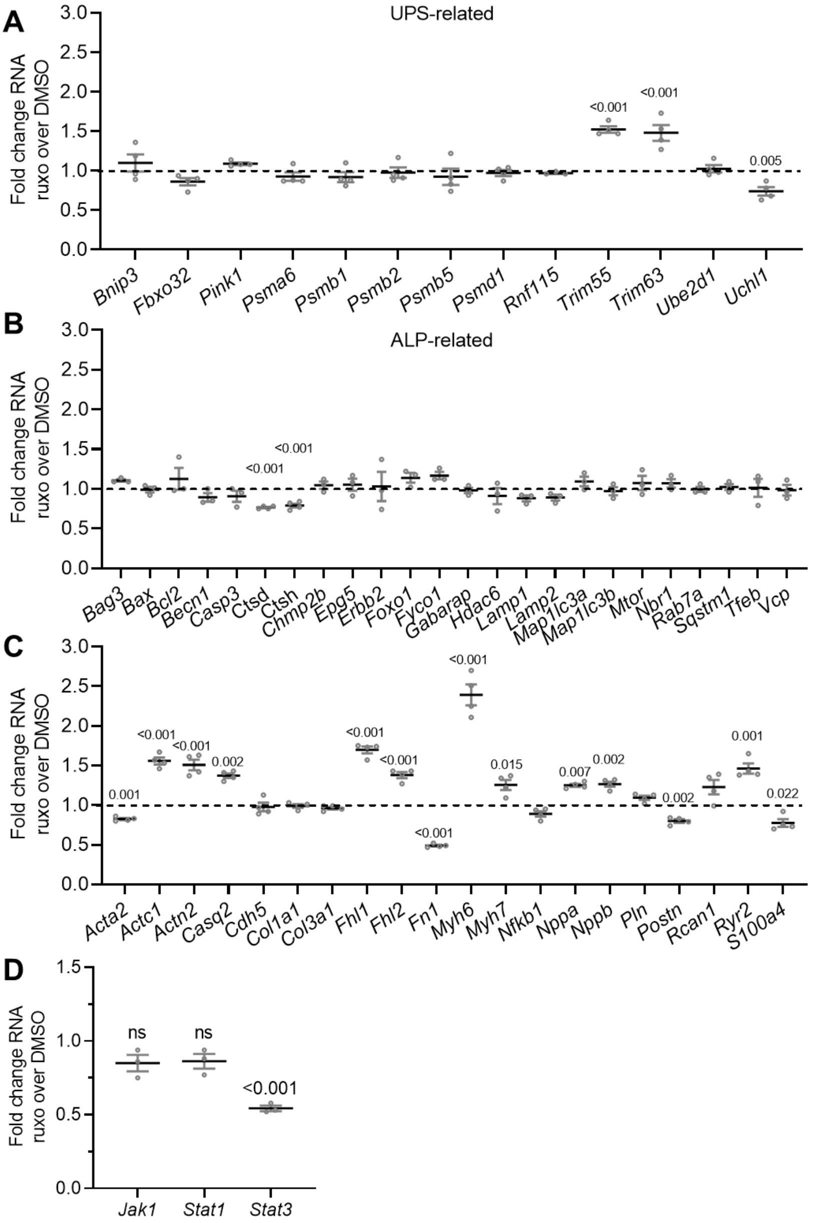


**Figure S7. NanoString RNA ratios of NRVMs treated with ruxolitinib to DMSO.** NRVMs were treated with 3 µM ruxolitinib or 0.1% DMSO and extracted RNAs were subjected to NanoString RNA analysis. **A**, UPS-related RNAs. **B**, ALP-related RNAs. **C**, Cardiomyocyte-relevant RNAs, **D**, JAK-STAT pathway-related RNAs. Data are presented as mean ± SEM, and p-values were obtained with the unpaired Student's t-test. Abbreviation: ns, non-significant.

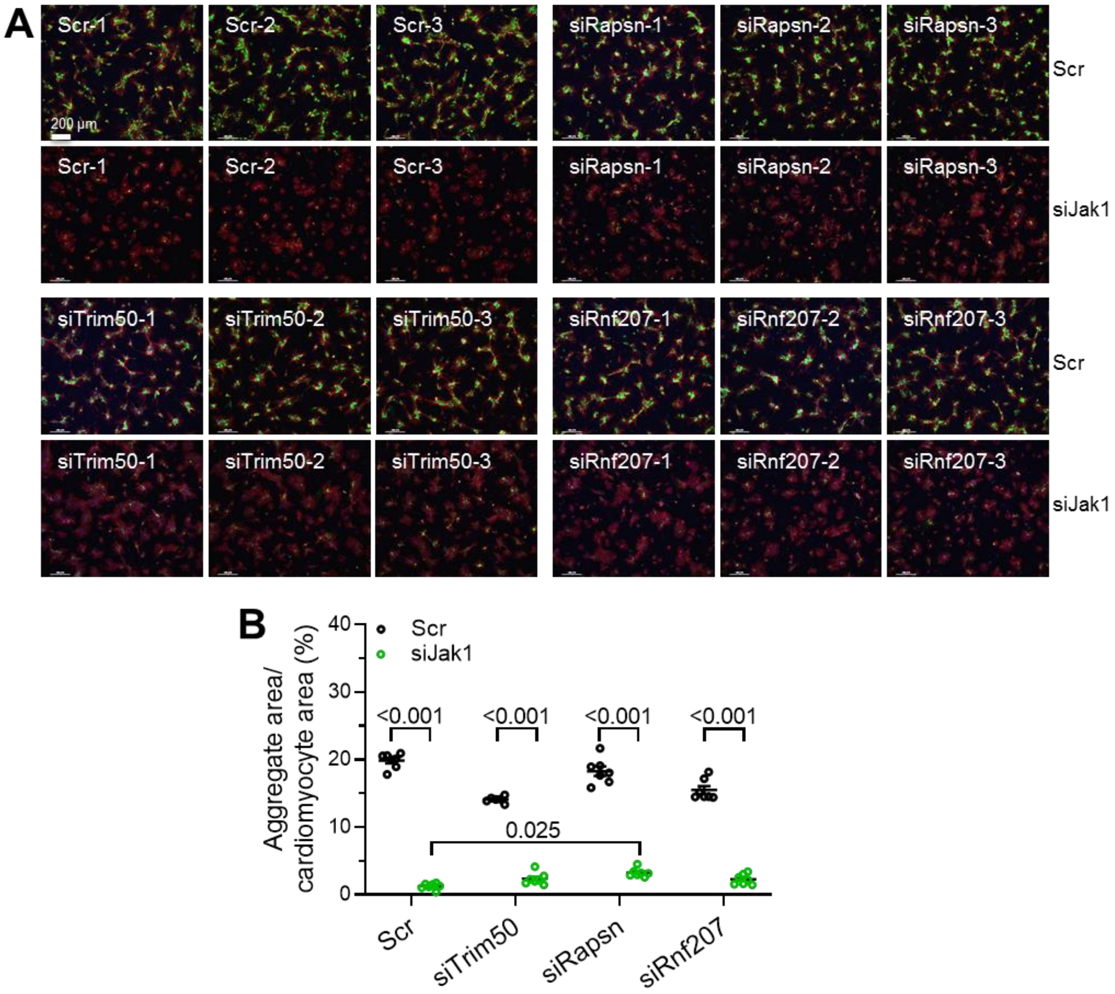


**Figure S8. Knockdown of E3 ligase *Trim50*, *Rapsn or Rnf207* has no major effect on CRYAB p.Arg120Gly aggregates after *Jak1* knockdown.** NRVMs transfected with 100 nM siTrim50, siRapsn, siRnf207 or scramble siRNA (scr), transduced with Ad5-CMV-CRYAB$^{R120G}$, treated with 3 µM ruxolitinib (ruxo) or DMSO and fixed after 5 days. **A**, Representative immunofluorescence images. Scale bar = 200 µm. Aggregates are depicted in green (CRYAB$^{R120G}$-GFP), cells in red (anti-cTnI) and nuclei in blue (DAPI). **B**, Quantification of aggregates in cardiomyocytes with NIS Elements software. Data are depicted as mean ± SEM, and p-values were obtained with the two-way ANOVA with Tukey's multiple comparisons post-hoc analysis. Dots represent images.

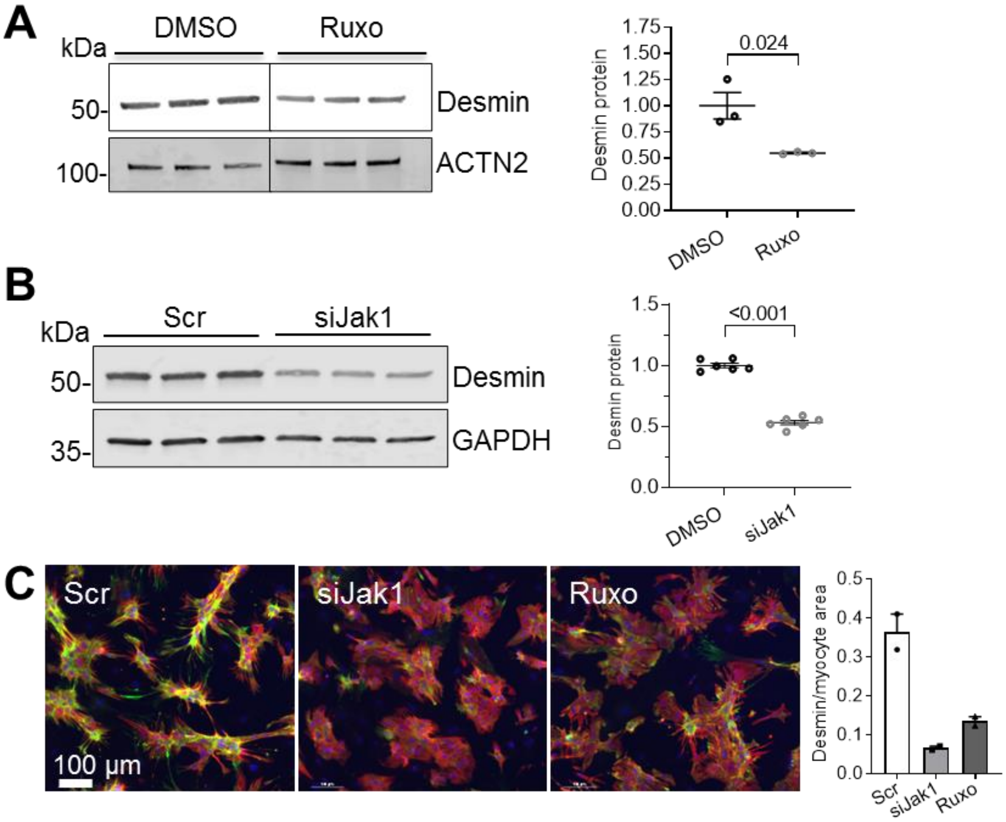


**Figure S9. Ruxolitinib or siJak1 treatment results in lower desmin protein levels. A,** Western blot of protein extracts from NRVMs treated with 3 µM ruxolitinib or 0.1% DMSO. Quantification of desmin protein level with ACTN2 as loading control. **B,** Western blot of protein extracts from NRVMs transfected with 100 nM siJak1 or scramble. Quantification of desmin protein level with GAPDH as loading control. **C,** Representative immunofluorescence images. Scale bar = 100 µm. Desmin is depicted in green (anti-desmin), cells in red (anti-cTnI) and nuclei in blue (DAPI). Quantification of aggregates in cardiomyocytes with NIS Elements software. Dots represent number of analyzed images.
